# Reconstructing Kinetic Models for Dynamical Studies of Metabolism using Generative Adversarial Networks

**DOI:** 10.1101/2022.01.06.475020

**Authors:** Subham Choudhury, Michael Moret, Pierre Salvy, Daniel Weilandt, Vassily Hatzimanikatis, Ljubisa Miskovic

## Abstract

Kinetic models of metabolic networks relate metabolic fluxes, metabolite concentrations, and enzyme levels through well-defined mechanistic relations rendering them an essential tool for systems biology studies aiming to capture and understand the behavior of living organisms. However, due to the lack of information about the kinetic properties of enzymes and the uncertainties associated with available experimental data, traditional kinetic modeling approaches often yield only a few or no kinetic models with desirable dynamical properties making the computational analysis unreliable and computationally inefficient. We present REKINDLE (REconstruction of KINetic models using Deep LEarning), a deep-learning-based framework for efficiently generating large-scale kinetic models with dynamic properties matching the ones observed in living organisms. We showcase REKINDLE’s efficiency and capabilities through three studies where we: (i) generate large populations of kinetic models that allow reliable *in silico* testing of hypotheses and systems biology designs, (ii) navigate the phenotypic space by leveraging the transfer learning capability of generative adversarial networks, demonstrating that the generators trained for one physiology can be fine-tuned for another physiology using a low amount of data, and (iii) expand upon existing datasets, making them amenable to thorough computational biology and data-science analyses. The results show that data-driven neural networks assimilate implicit kinetic knowledge and structure of metabolic networks and generate novel kinetic models with tailored properties and statistical diversity. We anticipate that our framework will advance our understanding of metabolism and accelerate future research in health, biotechnology, and systems and synthetic biology. REKINDLE is available as an open-access tool.

## Introduction

Recent technological progress in high-throughput measurement techniques has propelled discoveries in biology, biotechnology, and medicine, and allowed us to integrate multiple different data types into representations of cellular states and obtain deep insights into cellular physiology. Historically, researchers have used genome-scale models (mathematical descriptions of cellular metabolism), to associate experimentally observed biological data with cellular phenotype^1–3^. However, traditional genome-scale models cannot predict the dynamic cellular responses to internal or external stimuli because they lack information about metabolic regulation and enzyme kinetics^4,5^. Recently, the research community is shifting focus toward developing kinetic models of metabolism to further our understanding of cellular physiology^5^.

Kinetic models provide time-dependent predictions of cellular states and are amenable to sensitivity analysis, all of which allows to capture and model additional information about cellular metabolism compared to that obtained with traditional steady-state methods such as Flux Balance Analysis (FBA)^6–8^. However, the difficulty of acquiring the knowledge of (i) the exact mechanism of each reaction and (ii) the parameters of the said mechanisms hampers the building of kinetic models. In most of the current kinetic modeling methods^9–12^, the unknown reaction mechanisms are hypothesized or modeled by approximate reaction mechanisms^13,14^. The main challenge in obtaining the unknown parameters is qualitative and quantitative uncertainties intrinsic to biological systems. Due to the inherently under-determined nature of the mathematical equations describing the biological systems, it is frequent that the model can reproduce the experimental measurements for multiple rather than with a unique set of parameter values. The address these challenges, the researchers frequently choose Monte-Carlo sampling-based frameworks for the task at hand^15–19^. In these approaches, the space of admissible parameter values is first reduced by integrating the available experimental measurements and ensuring consistency with the physical and chemical laws. Then, the reduced solution space is sampled using Monte-Carlo techniques to extract a population of alternative sets of parameter values.

However, kinetic modeling frameworks that employ sampling to determine missing model parameters frequently produce large sub-populations of kinetic models that are inconsistent with the experimentally observed physiology. For instance, the constructed models can be locally unstable or display too fast or too slow time evolution of metabolite concentrations and metabolic fluxes compared to that of the experimental data (Fig.1), and hence, cannot be used for systems analysis and simulations. This entails a huge loss of computational resources and time, especially for cases where the incidence of sub-populations with desirable model properties is low. For example, we have observed in some studies that the rate of generation of locally stable large-scale kinetic models can be lower than 1%.^20^ These drawbacks become amplified with the increase in the size of the kinetic models because the size of sampling space explodes. As a result, finding regions in the parameter space that satisfy the desired properties and observed physiology becomes increasingly challenging.

**Fig. 1:**
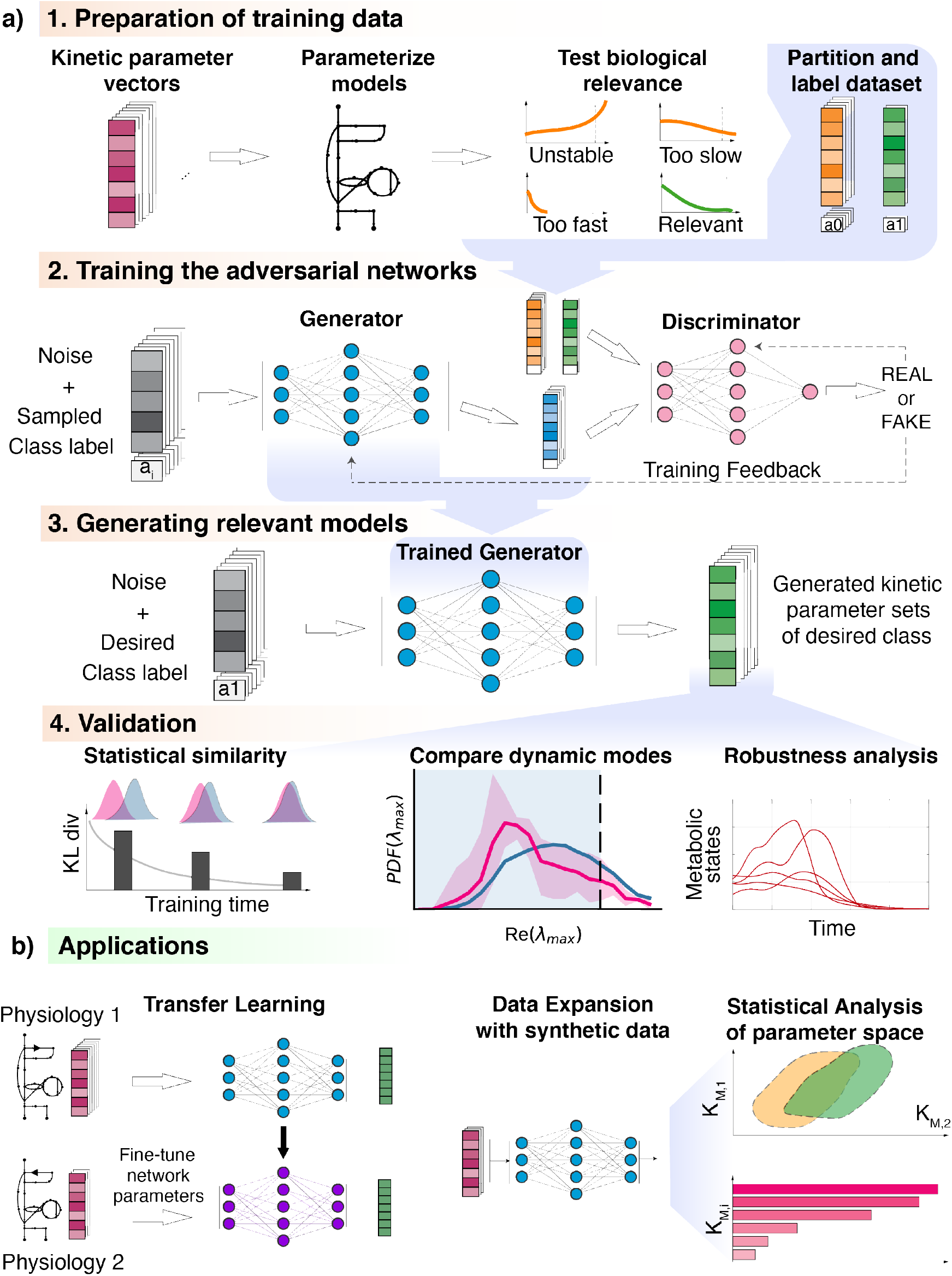
Overview of the REKINDLE framework and applications: **a) Framework:** Step 1 - Kinetic parameter sets are tested for pre-specified conditions (models that describe experimentally observed data and have appropriate dynamic properties) and are labelled and partitioned into data classes. Step 2 - REKINDLE employs Generative Adversarial Networks (GANs) to learn the distribution of labelled data obtained from the previous step. Step 3 - A trained generator from the GAN generates new kinetic parameters of models that satisfy pre-specified conditions. Step 4 **-** The generated dataset is subjected to statistical and validation tests for determining the satisfaction of the imposed conditions. **b) Applications:** REKINDLE uses the specifics of physiological and structural knowledge acquired by the GANs during training for extrapolating to other physiologies via transfer learning when training data is extremely limited (left). The REKINDLE-generated kinetic parameter sets are amenable to extensive and advanced statistical analysis allowing to reveal further insights about studied phenotypes (right).

These challenges call upon sophisticated statistical and machine learning methods that would facilitate the advances in this area of research. Sampling-based kinetic modeling frameworks lend themselves naturally to applications in deep learning because they can generate large sets of data required for the training of neural networks. The deep learning methods process the input raw data through several-layer-deep artificial neural networks and transform it into representations that are suitable for the interpretation and detection of patterns in the input.^21^ Among deep learning techniques, Generative Adversarial Networks (GANs) attracted considerable attention due to their capability to generate data.^22^

We present here REKINDLE (Reconstruction of KINetic Models using Deep LEarning), an unsupervised deep-learning-based method that leverages the generative ability of GANs to generate large populations of biologically relevant large-scale and genome-scale kinetic models. REKINDLE utilizes existing kinetic modeling frameworks to create the training data required to efficiently generate a large population of models with desired properties (Fig 1a). It significantly reduces the need for extensive computational resources and computation time required by the traditional kinetic modeling methods. For example, REKINDLE can be used to create large synthetic datasets within a matter of seconds on commonly used hardware. Importantly, we showcase REKINDLE’s ability to navigate through the physiological states of metabolism using Transfer Learning^23,24^ in the low data regime (Fig. 1b). REKINDLE’s departure from the traditional way of creating kinetic models paves a way for more comprehensive computational studies and advanced statistical analysis of metabolism.

### REKINDLE: A deep learning approach for generating biologically relevant large-scale kinetic models

The REKINDLE framework consists of four successive steps (Fig 1a). The input of REKINDLE are kinetic parameter sets obtained from kinetic modeling methods based on Monte-Carlo sampling^9,10,17,18,25^. The procedure starts by testing the biological relevancy of the kinetic parameter sets. Here, we consider that a kinetic parameter set is biologically relevant if the metabolic responses obtained from the kinetic model with this parameter set have experimentally observed dynamic responses. We then categorize the parameter sets into two classes, biologically relevant or not relevant (e.g., sets providing metabolic responses with a too slow, a too fast, or unstable dynamics), and label them accordingly (Fig 1a, Step 1). While in this study we use REKINDLE to generate kinetic models with biologically relevant dynamics, the framework allows imposing other bio-chemical properties or combinations of properties and physiological conditions to construct and label the dataset. A labeled dataset is then used for training the conditional GANs^26^ (Fig 1a, Step 2).

GANs consist of two feed-forward neural networks, the generator, and the discriminator. The goal of the training procedure is to obtain a good generator that generates kinetic models from a specific predefined class that are indistinguishable from the kinetic models of the same class in the training data (Methods). The discriminator gets trained by how well it could distinguish between the training data (real data) and the data provided by the generator (fake data) (Fig 1a, Step 3). The generator gets trained by how well it could deceive the discriminator into classifying its generated kinetic models as real. Both the discriminator and the generator get better at their tasks with more training steps. The generator gets better at deceiving the discriminator, and the discriminator gets better at classifying between training and generated data (Fig 1a, Step 2). The training continues until we reach equilibrium between the two neural networks, and no further improvement is possible.

Once the training is done, we validate the biological relevancy of the generated kinetic models via a series of tests (Fig 1a, Step 4). In this work, we performed three tests. We first tested the statistical similarity of the generated data with the training data by comparing their distributions in the parameter space. We then checked the distributions of the eigenvalues of the Jacobian (Methods). And their corresponding dominant time constants to verify if the generated parameter sets satisfy the desired time constraints. Finally, we tested the dynamic responses of the models to perturbations in the steady-state metabolic profile to the robustness of the generated parameter sets.

Further details about the REKINDLE framework can be found in the Methods section.

## Results

### REKINDLE generates large-scale kinetic models of *E. coli* metabolism

We showcase REKINDLE by generating biologically relevant kinetic models of the *E. coli* central carbon metabolism (Supplementary Fig. 1). The models are parameterized with 411 kinetic parameters (Methods). Thermodynamic-based flux analysis^27–29^ (TFA) performed on the model with the integrated experimental data from aerobic cultivation of wild-type *E. coli*^30^ indicated that two reactions, Transaldolase (TALA) and Isocitrate lyase (ICL), could operate in both forward and reverse directions, whereas the other reactions had unique directionalities (Table 1, Methods). This means that the study of *E. coli* metabolism in this condition requires generating four populations of kinetic models, with each population corresponding to a different combination of TALA and ICL directionalities. We enumerated these four cases as Physiology 1 – 4 (Table 1).

**Table 1:** Incidence of biologically relevant models generated with ORACLE (training data) and REKINDLE for four physiologies. The physiologies differ in the directions Transaldolase (TALA) and Isocitrate lyase (ICL) operate. TALA transforms Glyceraldehyde-3-phosphate to D-Fructose 6-phosphate in the Pentose Phosphate Pathway, and ICL converts Isocitrate to Succinate and Glyoxylate in the TCA cycle. The REKINDLE results represent the maximal incidence achieved for five repeats.

|  | Physiology1 | Physiology 2 | Physiology 3 | Physiology 4 |
| --- | --- | --- | --- | --- |
| Directionality of Reactions | $TALA \xrightarrow{\quad} ICL \xrightarrow{\quad}$ | $TALA \xrightarrow{\quad} ICL \xleftarrow{\quad}$ | $TALA \xleftarrow{\quad} ICL \xrightarrow{\quad}$ | $TALA \xleftarrow{\quad} ICL \xleftarrow{\quad}$ |
| ORACLE (training data) | 58.8% | 55.0% | 61.1% | 56.0% |
| REKINDLE | 97.7% | 97.3% | 99.3% | 100% |

While REKINDLE can use parameter sets of any kinetic modeling framework for the training, we employed the ORACLE framework^20,25,31–35^ implemented in the SKiMpy tool^36^ to generate a population of 80000 kinetic parameter sets for each physiology. The goal was to generate kinetic models that satisfy the observed steady-state and have dynamic responses that are faster than 6-7 minutes (which corresponds to 1/3 of the *E. coli* doubling time) and slower than 1 *μ*s. The kinetic models satisfying these two conditions can reliably reproduce experimentally measured metabolic responses in *E. coli* (Methods).

We analyzed the training data and observed that between 39% and 45% of models for the four physiologies have too slow dynamics (Table 1). This means that these models cannot be used to describe the dynamics of *E. coli* metabolism. We employed REKINDLE to improve the incidence of kinetic models that are consistent with the *E. coli* dynamics. To this end, we trained conditional GANs for a total of 1000 epochs with five statistical replicates for the four physiologies. The generator state was saved every 10 epochs and was tasked to conditionally generate 300 biologically relevant models. We quantified the dissimilarity in the distributions of the parameters generated by REKINDLE and the parameters generated by ORACLE (only the parameter sets corresponding to the biologically relevant dynamics) by calculating the Kullback-Leibler (KL) divergence (Methods) between the distributions of the two datasets. Here, we present the results for Physiology 1 (Table 1, Fig. 2), whereas the results for Physiology 2 – 4 can be found in the supplementary material (Supplementary Figure 3a-c, Supplementary 4a-c).

**Fig. 2:**
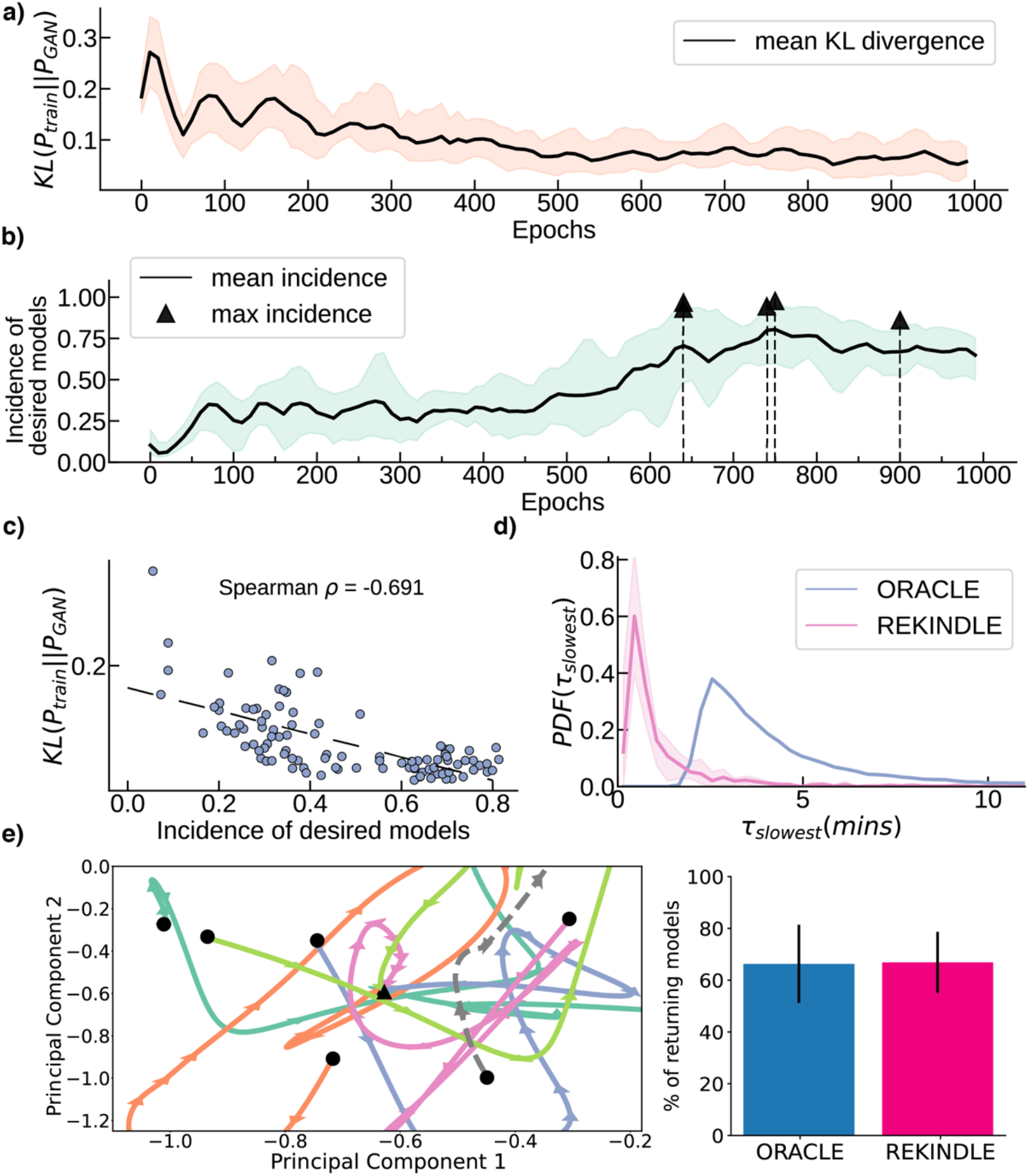
Generation and Validation of GAN generated kinetic models. **2a**. The similarity between the training data and generated data increases with training epochs as indicated by the decreasing Kullback-Leibler (KL) divergence between their distributions (black line) for 5 statistical replicates. Orange area indicates the KL divergence scores observed in five repeats. **2b**. The mean incidence of biologically relevant models in the generated data during training (black line). The black triangles indicate the maximum incidence attained for each repeat (Table 1). Green area indicates the incidence of biologically relevant models observed in five repeats. **2c**. Negative correlation of the mean incidence of relevant models with the KL divergence between the training and REKINDLE generated data. **2d**. REKINDLE shifts the dynamic responses of the generated data towards faster values, compared to the training data. Pink shaded area indicates overlap of distributions of *τ*_*slowest*_ in five repeats for the generators with the highest incidence of biologically relevant models. **2e. Perturbation Analysis:** (Left) Time-evolution of the randomly perturbed metabolic levels up to +-50% represented in the PCA space for five REKINDLE generated models. The black circles represent the initial perturbed state, the black arrows indicate the directionality of the trajectory in the PCA space, and the black triangle indicates the final, reference, state. The perturbed states either return to the reference steady-state (orange, blue, magenta, light green, dark green) or escape into a pathological state (gray dashed). (Right) Fraction of the perturbed models that return back to the reference steady-state for the training data produced by ORACLE and REKINDLE generated data.

We observed that the KL divergence decreases with training, meaning that the GAN learns the distribution of the kinetic parameters that correspond to biologically relevant dynamics (Fig 2a). The decrease in the KL divergence score also indicated that the GAN is not suffering from mode collapse, a common pathology in GANs where the generator maps the entire latent space to a small region in the feature space.^37^ In addition to monitoring the KL divergence during training, we also tested the generated models for biological relevancy using linear stability analysis of the resulting parameterized system of ODEs (Methods). The incidence of relevant models increased with the training epochs, reaching as high as 0.977 (97.7% of the generated models) for some repeats (Fig 2b, Table 1). Moreover, there was a strong negative correlation of ρ *=* -0.691 (Spearman correlation coefficient) between the number of relevant models at a given epoch and the KL divergence (Fig 2c). This indicates that the KL divergence is a good measure for assessing the quality of the training.

The training stabilized after ∼400 epochs (Supplementary Fig. 2a-d) and the discriminator accuracy sat around ∼50% (Supplementary Fig. 2a-d) suggesting that the GAN didn’t collapse and the generated models were not an artifact of a failed training process (Methods). It has to be noted that peak incidence of desired models occurred at a different number of epochs for different replicates (Fig 1b). In some cases, extending training to 1500 epochs was necessary to maximize incidence of desired models. Training beyond 1500 epochs showed negligible increase in performance. In some cases, training for too many epochs led to GAN collapse, i.e., the discriminator overpowered the generator.

### Validation of the REKINDLE generated models

We next set out to perform additional validation checks to determine the quality of the generated kinetic models. For all checks, we chose the generator with the highest incidence of desired models (Fig 2b) and used it to generate 10000 biologically relevant kinetic models.

In the first check, we verified how fast the dynamics of generated models is. To this end, we computed the distribution of the dominant characteristic time constant of the slowest dynamic response, *τ*, resulting from the generated kinetic parameter sets (Methods). We observed that the distribution of the time constants of the REKINDLE-generated models (Fig 2d, red) has shifted towards smaller values compared to the distribution of the time constants of the models from the training set (Fig 2d, blue). This meant that the dynamic responses of REKINDLE-generated models tend to have faster responses than the ones from the training set (Supplementary Note 1). That is, REKINDLE generates more physiologically relevant models than ORACLE. Indeed, the majority of the REKINDLE-generated models have the slowest dominant time constant of ∼1 minute (≪ 21 minutes, the doubling time of *E. coli*). In contrast, the majority of the models from the training set have a dominant time constant of 2.5 minutes with the distribution having a heavy tail towards larger time constants (Fig 2d, blue). We have observed similar results for the three remaining physiologies (Supplementary Figure 3a-c).

We next tested the robustness of the REKINDLE generated kinetic models by perturbing the steady-state and verifying if the perturbed system would evolve back to the steady-state (Fig 2e). We also performed the same test for the models from the training set and compared the results. For this purpose, we randomly chose 1000 biologically relevant parameter sets from both datasets (training and generated), and we perturbed the reference steady-state, *X*_*RSS*_, with random perturbations, Δ*X*, between 50%-200% of the reference steady state 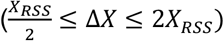. We repeated this procedure 100 times for each of the 1000 models. In the majority of cases, the system returns to within 1% of the steady-state at the *E. coli* doubling time, indicating that the kinetic models from the two datasets are locally stable around the reference steady-state and satisfy the imposed dynamic constraints. Indeed, the percentage of models coming back to the steady-state is comparable for both REKINDLE (66.85%) and ORACLE (66.31%) generated models (Fig 2e, right). We repeated the study for the remaining three physiologies, and we found that REKINDLE-generated models are consistently more robust than those generated by ORACLE. For example, for physiology 4, 83.79% of the REKINDLE generated models return to the steady-state, compared to 61.05% of the ORACLE generated models (Supplementary Figure 4a-c).

To visualize the time-evolution of the perturbed state of the kinetic models, we performed Principal Component Analysis (PCA) on the time-series data of the ODE solutions. The first two PCA components explained 97.17% of the total variance in the solutions (component 1: 85.21%, component 2: 11.95%). As an illustration, we plotted the first two PCA components of five randomly selected REKINDLE generated kinetic models that returned to the reference steady-state and one that escaped from it (Fig 1e, left).

These studies demonstrate that REKINDLE reliably generates kinetic models that are robust to perturbations and obey biologically relevant time constraints.

### Interpretability of the REKINDLE generated models

The efficiency of GANs in generating large populations of kinetic models lends itself well to a thorough statistical analysis of its outputs. REKINDLE captures the complex mapping between the kinetic parameters and the biologically relevant dynamics space, and interpreting its results will offer insights into the relevant kinetic parameters (features) that ensure the generated models satisfy imposed physio-biological conditions. To this end, we compared the population distributions of biologically relevant and irrelevant kinetic models generated by REKINDLE (2000 kinetic models from each category). We used the KL divergence to quantify the difference between the two distributions (Methods). The KL divergence score will allow us to identify the kinetic parameters that do and do not affect the biological relevancy of the overall kinetic model.

We observed that only a handful of parameters had a significant difference in the distributions between the two populations (Supplementary Figure 5a, b). This observation is consistent with previous studies showing that only a few kinetic parameters affect certain model properties,^38,39^ whereas the majority of parameters are sloppy^40^. We chose the top 10 parameters with the highest KL divergence score and inspected their distributions (Fig 3a). We observed that there is a clear bias in the distributions of the two populations (Fig 3a, right) suggesting that the values of these particular parameters are indeed affecting the biological relevance of the generated models.

**Fig. 3:**
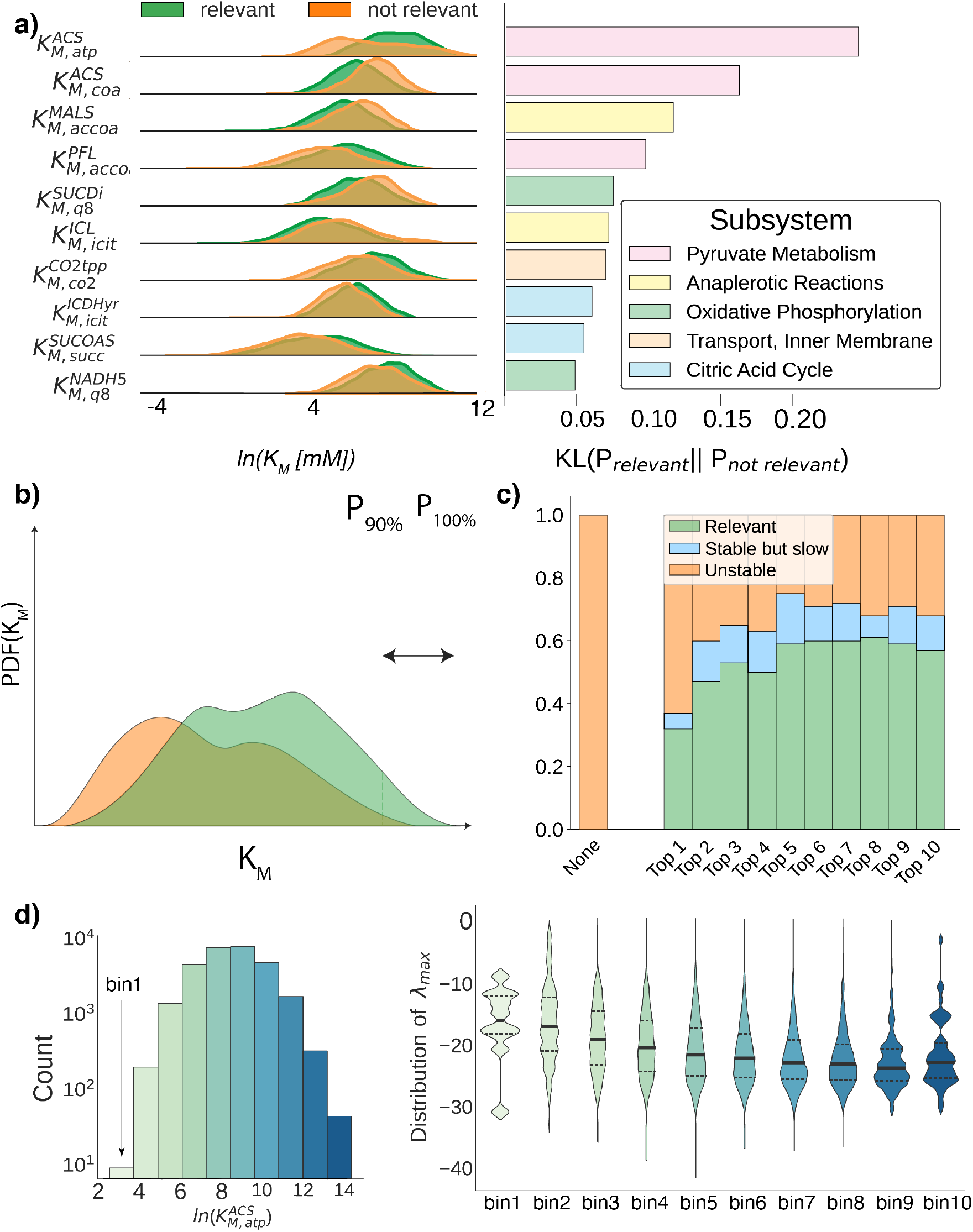
Interpretability of REKINDLE generated sets. **a) (Left)** Probability distribution comparison between relevant (green) and not relevant (orange) kinetic models for the top 10 ranked kinetic parameters (Supplementary Fig. 6, Supplementary Table S1). The parameters are ranked by the highest KL divergence between the two distributions. **(Right)** The respective magnitudes of KL divergence for the 10 parameters shown in (a). **b)** The parameters in (a) are constrained within the 90-th percentile and 100-th percentile (dashed vertical lines) of the relevant class (green) in the direction that favors biological relevancy (e.g., high values of 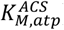). **c)** The fraction of locally unstable, stable, and relevant models as a result of constraining the parameters in (a) according to the constraints (b). We begin with 100 locally unstable models (orange bar, left) and gradually constrain the parameters in a cumulative manner, based on their ranks (right). Constraining the parameters stabilizes (blue) as well as rescues (green) the unstable models. We notice that there is negligible change in dynamics after the top 7 parameters have been constrained. **e)** (Left) 50,000 REKINDLE generated models were divided into 10 sub-populations (indicated by the different shades of the bins) based on their respective values of 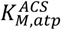. (Right) Maximum eigenvalue distributions of the Jacobian for the sub-populations. The thick black line indicates the mean of the distribution and the dashed lines indicates quartiles.

To test this hypothesis, we took a random population of 100 locally unstable models. Within this population, we constrained the highest-ranked parameter, 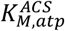 (Fig 3a, Supplementary Fig. 6, Supplementary Table S1), within its top 10 percentile values in the direction that favors relevancy, i.e., toward higher parameter values (Fig 3 b,c), and observed the dynamics of the resulting kinetic models. Remarkably, 29 of the resulting models became biologically relevant, meaning that this parameter strongly affects this property. We repeated this procedure and constrained the top 2 parameters, the top 3 parameters, and so on until we constrained all ten of them. In all cases, the constraints on the parameters resulted in a bigger portion of models becoming biologically relevant (Fig 3c). For example, when we constrained the top 7 parameters, we obtained ∼60% of models with biologically relevant dynamics and over 70% of locally stable models. We also observed that adding further constraints beyond the top 7 parameters did not significantly affect the dynamics, indicating that these parameters are sloppy.

Based on these results, we next hypothesized that a dependency of the system dynamics, quantified with the largest eigenvalue of the Jacobian, on the values of 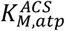 exists (Methods). To test this hypothesis, we generated 100000 biologically relevant models using the generator from above. We then split this dataset into ten different subsets according to the 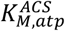 value, meaning that we had ten subpopulations of parameter sets with 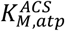 values ranging from low (Fig 3d, left, leftmost bar) to high (Fig 3d, left, rightmost). For each of the ten subpopulations, we computed the eigenvalue distribution (Fig 3d, right). As hypothesized, we observed that the mean of the eigenvalue distributions moves towards more negative values as the 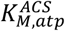 value increases (thick black line in Fig 3e, right). This means that the models will have faster dynamic responses for higher 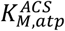 values. This is also consistent with the distributions of 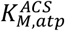 (Fig 4a, top parameter) showing that higher values of this parameter favor biological relevancy. We performed similar analysis for all of the parameters in Fig 4a. There were no such trends for the last few parameters confirming that these parameters are indeed sloppy (Supplementary Figure 7a,b).

**Fig. 4:**
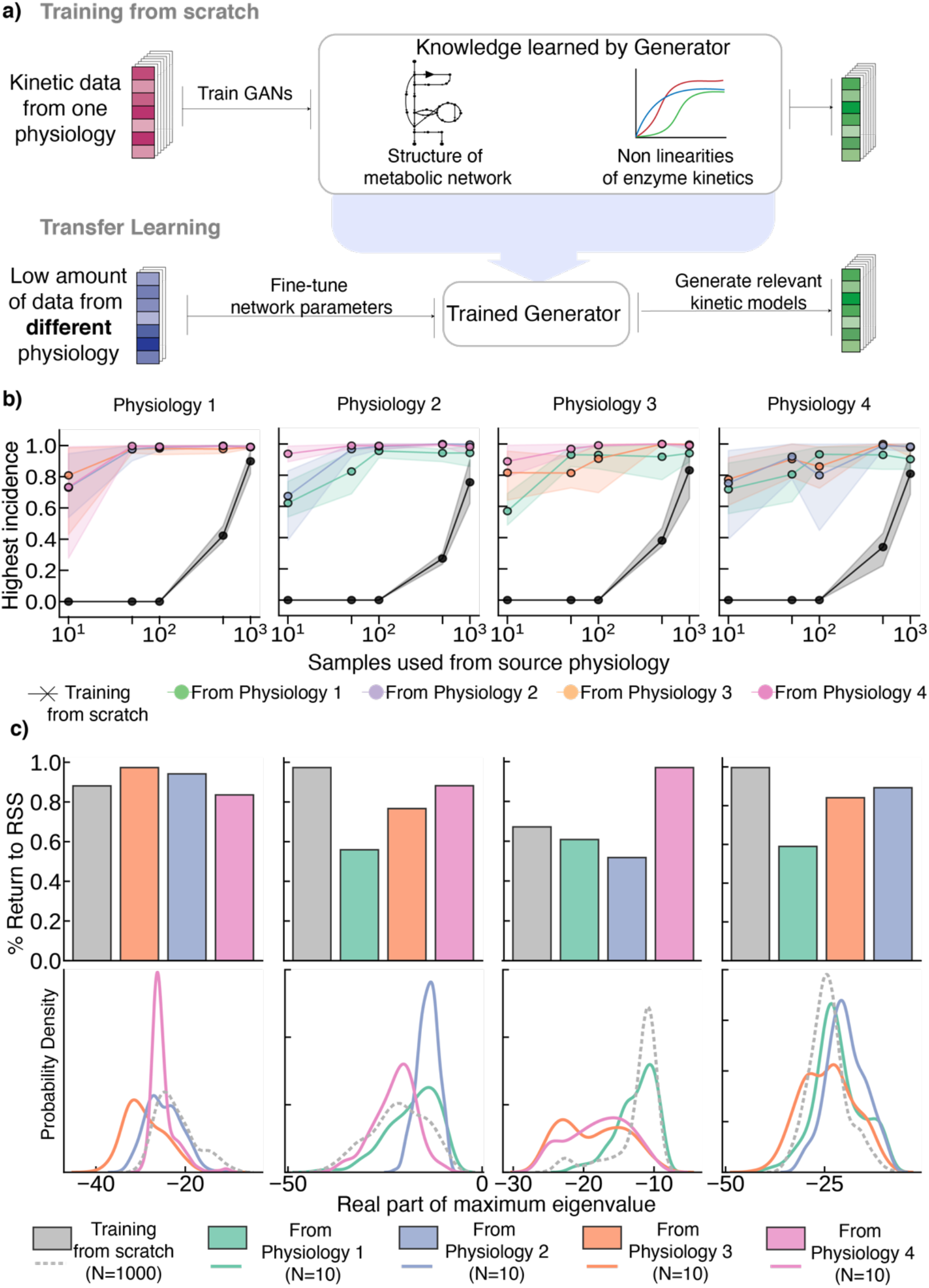
Extrapolation to multiple physiologies via Transfer Learning: **a)** A generator trained for one physiology learns the structure of the metabolic network and nonlinearities of enzymatic mechanisms allowing us to re-train it for another physiology with just a few parameter sets. **b)** Comparison of the incidence of biologically relevant models created with the generators trained from the scratch (gray crosses) and with transfer learning (colored circles) for four physiologies. For each physiology, we compare the training from the scratch and the transfer from the other three physiologies using different amounts of data (10, 50, 100, 500, and 1000 datasets). **c)** Validation of transfer learning. **Upper panel:** fraction of the perturbed models that return back to the reference steady-state for the models obtained from the scratch trained generators (gray) and models obtained from the transfer learning trained generators (green, transfer from Physiology 1, blue, transfer from Physiology 2, orange, transfer from Physiology 3, pink, transfer from Physiology 4). **Lower panel:** the probability density function of the real part of the maximum eigenvalue obtained for the populations of kinetic models obtained from the two types of generators (trained from the scratch and transfer learning). The color coding is consistent with the upper panel.

We repeated this study for the other three physiologies (Supplementary Fig. 8). Similar to Physiology 1, in Physiologies 2 – 4, the parameter group with the highest KL divergence scores had the highest impact on the biological relevance. While a high KL divergence score was an indication that a kinetic parameter was affecting the biological relevance, the parameter ranking on this score was not always coinciding with the most efficient conversion of biologically irrelevant models to relevancy. For example, we observed for Physiology 3 that biologically irrelevant models became relevant only after constraining the first five parameters (Supplementary Fig. 8), suggesting that the 5^th^ ranked parameter was the most significant for biological relevancy.

These results suggest that GANs trained in REKINDLE are capable of implicit model reduction and distinguishing between features (kinetic parameters) that directly affect desired properties, which in our case are biologically relevant dynamics properties, from features that do not. To confirm this, we performed unsupervised dimension reduction with UMAP^41^ on REKINDLE generated set and the training set for Physiology 1, first using all the parameters and then using only the top 5 parameters from Fig. 3A (Supplementary Fig. 10a, b). In the former case, we observed that the generated set and the training set are disjoint, while in the latter case both sets share the same space. This shows that the GANs distill important information primarily learning the distributions of significant kinetic parameters and giving less importance to the parameters that do not affect the desired kinetic property.

### Extrapolation to other physiologies using transfer learning in low data regime

Adding to the requirements on biologically relevant time scales, most kinetic modeling methods manage to find only a few parameter sets consistent with the experiments due to the highly non-linear and stiff nature of the ODEs in metabolic networks and high-dimensional parameter space. Even when we can generate large parameter sets, the number of valid parameter sets usually significantly diminishes after conducting consistency checks and pruning to accommodate experimental observations.^5^ Therefore, for comprehensive analyses of metabolic networks, we need to create large populations of parameter sets. However, there is a trade-off between generating large datasets and computational requirements, and, depending on the efficiency of employed methods, this results in the limited scope of the majority of studies.

REKINDLE addresses this issue by using the extrapolation abilities of GANs via transfer learning^42^. In transfer learning, a machine learning model trained for one task is used to perform a related but different task. Here, we took a generator that has been trained for one physiology and fine-tuned it for another physiology using a considerably smaller set of training models (Fig 4a). To illustrate benefits of this approach, we considered four different physiologies (Table 1, and Methods). We started by taking the generator previously trained for Physiology 1 and used it to retrain GANs for the 3 remaining physiologies using 10, 50, 100, 500, and 1000 training kinetic models (computed by ORACLE). For each case, we trained GANs for a total of 300 epochs and repeated the training 5 times for every physiology with a randomly weighed discriminator (Methods, Supplementary Fig. 11a, b, c). Remarkably, the transfer learning with only 10 parameter sets provided considerably high incidence of biologically relevant models of Physiologies 2-4 (Fig 4c, blue, orange, and violet circles). Indeed, the incidence ranged from 72% (for the transfer from Physiology 1 to Physiology 3 and 4) to 82% (from Physiology 1 to Physiology 2). More strikingly, the transfer attained the incidence of 100% for all three physiologies already as of 50 parameter sets.

For objective comparison, we then trained GANs from scratch for Physiology 2 - 4 using 10, 50, 100, 500, and 1000 training parameter sets for a total of 1000 epochs in each training. Despite the shorter training (300 versus 1000 epochs), the transfer learning significantly outperforms training from scratch (Fig 4b). Training from scratch reaches a comparable incidence of relevant models to the one of transfer learning only for around 1000 parameter sets. Training from scratch completely fails when the number of samples is below 500 as the discriminator easily overpowers the generator43 (Fig 4b). The minimum amount of data required to train a randomly initialized GAN for reliably generating relevant models depends on the sizes of the used neural networks and the studied metabolic system. Determining the minimal amount of data for training as a function of the size of the studied metabolic system remains an open problem.

In the same fashion, we performed the transfer learning from Physiology 2, 3 and 4 to the other three physiologies (i.e., from Physiology 2 to Physiology 1, 3, and 4, etc.). As expected, the transfer learning required significantly less amount of data and training epochs compared to when training is done from scratch (Fig 4b). The transferred generators displayed good extrapolation results even when provided with only 10 models from the target physiology with the average incidence of biologically relevant models being between 0.6 to 0.8. The GANs for Physiologies 1 - 3 trained by transfer learning from Physiology 4 performed exceptionally well, with the incidence of feasible models reaching ∼100% for 100 training sets (Fig 4b, pink circles).

We further compared the features of the kinetic models obtained from the generators trained from scratch and the ones generated via transfer learning (Fig 4c). Similar to the study from Fig 2e, we first investigated how many models will evolve back to the reference steady-state when their metabolic state is perturbed. The results showed that the kinetic models generated via transfer learning have similar robustness properties as those from GANs trained from scratch despite using the much smaller dataset, i.e., only 10 parameter sets (Fig 4c, upper panel). We then set off to compare the kinetic parameter distribution of the two classes of models. A narrow parameter distribution could indicate that the generated models stem from a constrained region in the space and that generators are not producing diverse kinetic models. Instead of comparing the distributions of individual kinetic parameters, we used the distribution of the largest negative eigenvalue of the generated kinetic models as a meta-measure for the spread of parameter distributions (Fig 4c, lower panel). We observed that the models generated via transfer learning have a well-spread distribution comparable to one of the models generated via learning from scratch.

We conclude that transfer learning with only a few kinetic parameter sets allows the generation of kinetic models that possess the desired property of biological relevancy, that are robust and parametrically diverse. We anticipate that this approach could help to derive new ways for high throughput analysis of metabolic networks.

## Discussion and Conclusions

The scarceness of experimentally verified information about intracellular metabolic fluxes, metabolite concentration, and kinetic properties of enzymes leads to a highly undetermined system with multiple models capable of capturing the experimental data. Due to the requirement of intense computational resources aimed to quantify the involved uncertainties, researchers often end up using only one out of the many alternative solutions, leading to unreliable analysis and misguided predictions about metabolic behavior of cellular organisms. This is the single most important reason for the limited use of kinetic models in the studies of metabolic systems despite their wide-acknowledged capabilities. REKINDLE offers a new, highly efficient way of creating large-scale kinetic models, thus enabling an unprecedented level of comprehensiveness for analyzing these networks and offering a much broader scope of applicability of kinetic models. Moreover, REKINDLE offers a highly efficient way for sampling the parameter space. In general, sampling of nonlinear parameter spaces has emerged as a standard method in addressing under-determinedness in computational physics, biology, and chemistry^44,45^.

The proof-of-concept applications presented here demonstrate REKINDLE’s ability to learn the mechanistic structure of the metabolic networks and to stratify kinetic parameter subspaces corresponding to relevant model properties. By learning a map between the complex high dimensional space of kinetic parameters and a less complex embedding vector (Gaussian noise used as generator seed), the GANs essentially augments (i) the efficiency at which we can create models corresponding to our specified criteria and (ii) the information available to partition the parameter space according to our criteria. Consequently, REKINDLE is several orders of magnitudes faster in generation, compared to traditional methods, owing to its usage of easily available, widely used and highly abstracted linear-algebra libraries (Methods). When generators are trained from scratch, REKINDLE requires a minimum of 1000 data points to reliably reach high incidence of relevant models (Fig 4b). This corresponds to a training time of ∼15-20 minutes (Table 2). A trained generator in REKINDLE generates 1 million models in ∼18 seconds. In comparison, ORACLE, currently one of the most efficient kinetic modeling frameworks, accomplishes the same task in 18-24 hours with identical computational power and on the same operational system (Table 2). The reduction in generation time is even more pronounced when models are generated through transfer learning due to the low amount of data being used for training.

**Table 2:** Comparison of computational time between ORACLE, REKINDLE, and REKINDLE with transfer learning (REKINDLE-TL). The generation time for 1 million biologically-relevant kinetic models reduces by more than six thousand times when REKINDLE is used instead of ORACLE (last column). In total, when including the generation of training data and training time, REKINDLE and REKINDLE-TL are more efficient than ORACLE by more than 60 and 600 times, respectively, for generating 1 million relevant models.

|  | Generation of Training data | Training Time | Generation of 1 million kinetic models | Generation of 1 million **relevant** kinetic models |
| --- | --- | --- | --- | --- |
| ORACLE | - | - | ~ 18-24 hours | ~ 36-48 hours |
| REKINDLE | ~ 15-20 mins (1000 models) | ~ 15 mins (1000 models) | ~ 15-20 seconds | ~ 15-20 seconds |
| REKINDLE-TL | ~ 5 seconds (10 models) | ~ 2-3 mins (10 models) | ~ 15-20 seconds | ~ 15-20 seconds |

Once a good generator is trained for the target physiology via transfer learning using low number of data samples, the generator can be used to expand upon traditionally small datasets with newly generated synthetic dataset, within seconds. Such expanded datasets are amenable for traditional statistical analyses to gain further knowledge about the system being studied. In contrast, this kind of analysis is not possible with small datasets. This offers to REKINDLE a significant advantage in scope of applications and comprehensiveness over traditional methods for generating large-scale kinetic models.

As is the case for many deep-learning-empowered frameworks, finding the optimum architecture of the networks used in REKINDLE concerning the size of the studied metabolic network remains an open problem. Addressing this question will open up the possibility of constructing highly curated libraries of ‘off-the-shelf’ networks that have been pre-trained using commonly used standard metabolic models. This will provide researchers in academia and industry with a repository to apply this framework to different physiologies and different types of studies and applications ranging from biotechnology to medicine.

In summary, we present a framework to generate kinetic models leveraging the power of deep learning while at the same time retaining the convenience of traditional analyses, where researchers can analyze the structural dependencies, correlations, and feedbacks within the metabolic network. The open-access code of REKINDLE will allow a broad population of experimentalists and modelers to couple this framework with experimental methods and benefit from synergistic approaches for the analysis and metabolic interventions of studied organisms.

## Methods

### Kinetic models of wild-type *E. coli*

Kinetic nonlinear models used in this study represent the central carbon metabolism of wild-type *E. coli*. They are based on the model published by Varma and Palsson^30^ and studied extensively using the SKimPy toolbox^36^ (Supplementary Fig. 1). The kinetic models consist of 64 metabolites, distributed over cytosol and extracellular space, and 65 reactions including 16 transport reactions. Out of the 64 metabolites in the model, 15 metabolites are boundary metabolites, i.e., they are localized in the extracellular space. The remaining 49 metabolites are localized in the cytosol and their mass balances are described with the ordinary differential equations (ODEs). After assigning each reaction to a kinetic mechanism based on their stoichiometry, we parameterized this system of ODEs with 411 kinetic parameters.

### ORACLE framework and generation of the training set

The ORACLE framework consists of a set of computational procedures that allow us to build a population of large-scale kinetic models that account for the uncertain and scarce information about the kinetic properties of enzymes.^20,25,31–35,46^. The idea behind ORACLE is to reduce the feasible space of the kinetic parameters through the integration of available experimental measurements, omics data, and thermodynamic constraints and then employ Monte-Carlo sampling techniques to determine unknown parameters^15,47^. ORACLE builds the kinetic models around the thermodynamically consistent reference steady-state fluxes and metabolite concentrations^31,48,49^. Instead of directly sampling the kinetic parameters such as Michaelis-Menten constant and inhibitory constants, we sample the enzyme saturation in the following form:^42^

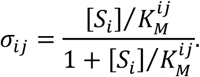

Then, we back-calculate the values for the kinetic parameters, 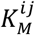, using the knowledge of the steady-state concentrations, [*S*_*i*_], and the enzyme saturation, *σ*_*ij*_. Once we know the kinetic parameters, 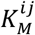, the equilibrium constants, 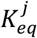, obtained when calculating the thermodynamically consistent steady-state, and steady-state flux, *v*_*j*_, we can calculate the maximal velocities, 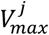, by substituting these quantities in the rate expressions. This way, the kinetic model is completely parameterized.

### Determining biological relevancy and partition of dataset into sub-populations

To guarantee that the generated kinetic models are capable of reliably describing metabolic processes, we have to ensure that they are locally stable and have biologically relevant dynamics. To this end, we compute the Jacobian of the dynamic system^47^. The sign of the eigenvalues of the Jacobian gives us information on whether or not the generated models are locally stable. For a model, if all eigenvalues are negative, then the model is locally stable. Otherwise, if any of the eigenvalues are positive, the model is unstable. Moreover, for a locally stable system, we can calculate from the Jacobian the characteristic time constant of the linearized system as the inverse of the real part of the largest eigenvalue of the Jacobian. Models having the largest eigenvalues with a negative real part close to zero will yield slow dynamics that can be slower than dynamics of metabolic processes observed experimentally.

Therefore, to ensure that the characteristic time constants of generated models is biologically relevant, we considered that this quantity is at least three times faster than the doubling time of *E. coli* (∼21 minutes). Here, we assume that the metabolic responses are aperiodic and that after three characteristic times the responses reach ∼95% of the steady-state. Therefore, to ensure this, we impose a strict upper bound of Re(λ_i_) < −9 on the real part of the Jacobian eigenvalues, λ_i_. The dominant time constant for this eigenvalue is then calculated as 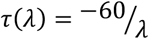 (in minutes). All kinetic parameter sets that obey this constraint are labelled as biologically relevant, otherwise they are labelled irrelevant.

### Data preprocessing

After labelling, the dataset was log transformed as the kinetic parameters spanned several orders of magnitudes. The dataset was then split into training and test sets with the ratio 9:1 which resulted in 72000 models that were used for training the cGAN. Moreover, only the concentration associated parameters, 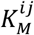, were included as the training features, because the scaling coefficients, 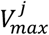, can be calculated back once the 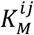 and steady state concentration and flux profiles are already known. Thus, after eliminating 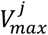, the training dataset consisted of 259 features in total.

It should be noted that both classes of models in the training data, biologically feasible and infeasible, share significant overlap in the kinetic parameter space and cannot be independently visualized by low-order dimension reduction techniques like principal component analysis (PCA)^50^, t-SNE^51^, and UMAP (Supplementary Fig. 10)

### Perturbation analysis of kinetic models

We randomly choose 1000 biologically relevant kinetic parameter sets from both ORACLE sampled dataset and REKINDLE generated dataset for any given physiology. We parameterize the system of ODEs describing the metabolite concentrations using REKINDLE or ORACLE generated kinetic parameter sets. Then, for each parameterized system we randomly perturb the reference steady state metabolite concentration (*X*_*ref*_) and flux profile of the model up to ±50%. We next integrate the ODEs using this perturbed state *X*’ as the initial condition *X*(*t* = 0) = *X*_0_. To quantify if a model came back to the reference steady state, we monitor the L_2_ norm of the metabolite concentrations at a given point of time *X*(*t*) and the reference concentration *X*_0_. If the metabolite concentration at 21 mins (doubling time of *E. coli*) is less than 1% of the reference steady state we classify the model as having returned back to the steady state, i.e., we test,

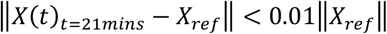

We repeat this process 10 times for each parameter set with a random perturbation each time (within ±50%).

### Kullback-Leibler Divergence / Relative Entropy

For two separate probability distributions P(x) and Q(x) over the same random variable x, we can measure how different these distributions are using the Kullback-Leibler divergence^52^ from Q(x) to P(x) is formulated as,

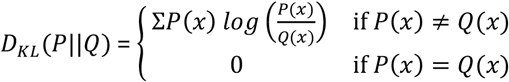

### Spearman correlation coefficient

For two random variables *X* and *Y*, we compute the Spearman correlation coefficient as

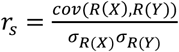

where *R*(*X*) and *R*(*Y*) are the ranks of *X* and *Y*, respectively, *cov*(*R*(*X*), *R*(*Y*)) the covariance of the rank variables, and *σ*_*R(X)*_ and *σ*_*R(Y)*_ are the standard deviations of the rank variables.

### GAN training

In Generative Adversarial Networks (GANs), two neural networks, the generator that we train to generate new data and the discriminator that tries to distinguish generated new data from real data, are pitted against each other in a zero-sum game. The end goal of this game is to obtain the generator that generates new data of such a quality that the discriminator cannot distinguish it from real data (Fig 1a, Step 3). We train the generator and discriminator networks in turns. For training the discriminator, we freeze the generator by fixing its network weights. Then, we alternate the training by freezing the discriminator and train the generator. In the first part of each learning step, we provide the discriminator with (i) a random batch of kinetic parameter sets from the training data with labels indicating the class of models and (ii) a batch of kinetic parameter sets that have been generated by the Generator (fake data). The discriminator then classifies the models it is presented with as real (coming from the training set) or fake (coming from the generator). In the second part of a learning step, the discriminator is frozen and the generator generates a batch of fake kinetic parameter sets using as inputs (i) random Gaussian noise and (ii) sampled labels. The discriminator and the generator improve their performance with training. The generator improves it capability to create “real-like” data, and the discriminator better distinguishes between the “real-like” and real data. The training stops when there is no further improvement.

### GAN implementation

All software programs were implemented in Python (v3.8.3) using Keras (https://keras.io/, v2.4.3) with Tensorflow^53^ GPU backend (www.tensorflow.org, v2.3.0). The GANs were implemented as conditional GANs. The discriminator network was composed of 3 layers that have a total of 18319 parameters: layer 1, Dense with 32 units, Dropout (0.5);layer 2, Dense with 64 units, Dropout (0.5); layer 3, Dense with 128 units, Dropout (0.5). The generator network was composed of 3 layers that have a total of 315779 parameters: layer 1, Dense with 128 units, BatchNormalization, Dropout (0.5); layer 2, Dense with 256 units, BatchNormalization, Dropout (0.5); layer 3, Dense with 512 units, BatchNormalization, Dropout (0.5). We used the Binary Cross-Entropy loss and the Adam optimizer^54^ with a learning rate of 0.0002 for training both the networks. We used batch sizes of 2, 5, 10, 20, and 50 when training with training set sizes of 10, 50, 100, 500, and 1000, respectively. We trained the GANs from scratch over each physiology for 1000 epochs (one epoch is defined as one pass over all of the training data) which took approximately 40 minutes for training over 72000 models on a single Nvidia Titan XP GPU with 12 GB of memory. Training was repeated 5 times for each physiological condition with randomly initialized generator and discriminator networks (20 total trainings).

In all studies, we considered that training has failed if the discriminator accuracy is consistently greater than 90% for the last 200 epochs.

### Generation of biologically irrelevant kinetic models using trained generators

To study the statistical differences in the distributions of the biologically relevant and non-relevant kinetic models we generated two populations (2000 models) of each class using a trained generator. As most of the trained generators do not have 100% incidence of biologically relevant models (Table 1), they have a small incidence of non-relevant kinetic models even when conditionally generating relevant models. We use this to create a population of non-relevant kinetic models, by continuously generating relevant models using the respective conditional label until we obtain a population of 2000 non-relevant models. Alternatively, we can also generate non-relevant kinetic models by directly using the appropriate conditional label with the generator seed during generation. However, when we subjected the population generated in this manner to statistical analysis (calculating the KL divergences between the individual kinetic parameters) we failed to retrieve the significant parameters. We hypothesize that this due to the absence of significant kinetic parameters that determine the local instability of a kinetic model (Supplementary Note 1).

### Transfer Learning on multiple physiologies

We trained GANs from scratch for each physiology using 72,000 samples. We then saved the generator state for the generator that had the highest incidence of relevant models. Then we re-trained this generator in a GAN setting with a randomly weighted discriminator using a low amount of data (10, 50, 100, 500 and 1000) from the target physiology on which extrapolation is desired, for 300 epochs. For the instances using 500 and 1000 samples the learning rate was changed from 0.0002 to 0.001. For the rest of the cases, we trained with the same hyperparameters as discussed in the previous section. During training, we generated 1000 biologically relevant models every 10 epochs and calculated the eigenvalues of the Jacobian, as previously, to monitor the quality of training and the ability of the generator to generate biologically relevant models from the target physiology. We repeated the training 5 times for each transfer (Physiology a to Physiology b) with the same pre-trained generator (on Physiology a) but with randomly weighted discriminator each time. We noted the highest incidence of relevant models during training save the corresponding generator state for each instance of training. We repeated this entire procedure (transferring generator, retraining GAN, generation of relevant models and validation using eigenvalues of Jacobian) for different amounts of data from the target physiologies (10, 50, 100, 500 and 1000 samples). Total training counts: 4 (total physiologies) × 3 (target physiologies) × 5 (repeats)× 5 (amount of data used) = 300. The resulting highest incidences of biologically relevant models from the target physiology as a function of the amount of data samples used is summarized in Fig 4b.

## Code and data availability

All code and data used in our study is publicly available. Data is available from the Zenodo repository (https://zenodo.org/record/5803120 and the links therein). A TensorFlow implementation of the REKINDLE workflow is available at https://github.com/EPFL-LCSB/rekindle and https://gitlab.com/EPFL-LCSB/rekindle. The ORACLE framework is implemented in the SKimPy (Symbolic Kinetic models in Python)^36^ toolbox, available at https://github.com/EPFL-LCSB/skimpy.

## Acknowledgements

This work was supported by funding from the Swiss National Science Foundation grant 315230_163423, the European Union’s Horizon 2020 research and innovation programme under grant agreements 722287 (Marie Skłodowska-Curie) and 814408, Swedish Research Council Vetenskapsradet grant 2016-06160, and the Ecole Polytechnique Fédérale de Lausanne (EPFL). We gratefully acknowledge the support of NVIDIA Corporation with the donation of two Titan XP GPUs used for this research.

## Supplementary Information

**Supplementary Figure 1:** Escher map of the E. coli central carbon metabolism network.

**Supplementary Figure 2:** Discriminator loss, generator loss and discriminator accuracy when training from scratch for (a) Physiology 1 (b) Physiology 2 (c) Physiology 3 (d) Physiology 4

**Supplementary Figure 3:** Generation and validation of REKINDLE generated datasets for (a) Physiology 2 (b) Physiology 3 (c) Physiology 4

**Supplementary Figure 4:** Perturbation analysis of REKINDLE generated datasets for (a) Physiology 2 (b) Physiology 3 (c) Physiology 4

**Supplementary Figure 5:** KL divergence scores between the distributions of individual kinetic parameters in the relevant and non-relevant classes of REKINDLE generated kinetic parameter sets for Physiology 1

**Supplementary Figure 6:** The top 7 kinetic parameters from Fig 3a are visualized in the metabolic network.

**Supplementary Figure 7:** Distribution of the eigenvalues of the Jacobian of sub-populations based on the values of the 2 lowest ranked parameters in Fig 3a.

**Supplementary Figure 8:** Highest ranked parameters based on KL divergence scores between relevant and non-relevant REKINDLE generated kinetic parameter sets, the respective metabolic subsystems they belong to and the effect of constraining them for Physiology 2, 3 and 4.

**Supplementary Figure 9:** Visualization of the kinetic parameter space of the training data for Physiology 1 using (a) PCA (b) UMAP (c) tSNE.

**Supplementary Figure 10:** Visualization of the kinetic parameter space of the training data and REKINDLE generated data for Physiology 1 using UMAP (a) for all the kinetic parameters (b) top 5 parameters from Fig 3a.

**Supplementary Figure 11:** (a) Discriminator loss and generator loss for transfer learning from Physiology 4 to 1 using 10, 50, 100, 500 and 1000 samples from Physiology 1. (b) Discriminator Accuracy for the same specifications as (a) (c) Incidence of biologically relevant models for the same specifications as (a).

**Supplementary Figure 12:** Differences in distributions of individual kinetic parameters (y-axis) using KL divergence between 16 discretized sub-populations based on eigenvalue space (x-axis) near the class boundary i.e., Re(λ_max_) = −9 for (a) Physiology 1 (b) Physiology 2 (c) Physiology 3 (d) Physiology 4.

**Supplementary Figure 13:** Differences in distributions of individual kinetic parameters (y-axis) using KL divergence between 16 discretized sub-populations based on eigenvalue space (x-axis) near the class boundary i.e., Re(λ_max_) = −9 for (a) Physiology 1 (b) Physiology 2 (c) Physiology 3 (d) Physiology 4. Here the KL divergence for each sub-population is normalized between 0 and 1.

**Supplementary Note 1:** Generation of biologically irrelevant models using REKINDLE.

## Abbreviations

GAN: Generative Adversarial Networks
ORACLE: Optimization and Risk Analysis of Complex Living Entities
TFA: Thermodynamics-based Flux Balance Analysis
GEM: GEnome-scale Model
REKINDLE: REconstruction of KINetic models of metabolism using Deep LEarning
FBA: Flux Balance Analysis
ODE: Ordinary Differential Equations

## References

1. Förster, J., Famili, I., Fu, P., Palsson, B. Ø. & Nielsen, J. Genome-Scale Reconstruction of the Saccharomyces cerevisiae Metabolic Network. Genome Res 13, 244–253 (2003).

2. Duarte, N. C. et al. Global reconstruction of the human metabolic network based on genomic and bibliomic data. Proc National Acad Sci 104, 1777–1782 (2007).

3. Seif, Y. & Palsson, B. Ø. Path to improving the life cycle and quality of genome-scale models of metabolism. Cell Syst 12, 842–859 (2021).

4. Almquist, J., Cvijovic, M., Hatzimanikatis, V., Nielsen, J. & Jirstrand, M. Kinetic models in industrial biotechnology – Improving cell factory performance. Metab Eng 24, 38–60 (2014).

5. Miskovic, L., Tokic, M., Fengos, G. & Hatzimanikatis, V. Rites of passage: requirements and standards for building kinetic models of metabolic phenotypes. Curr Opin Biotech 36, 146–153 (2015).

6. Orth, J. D., Thiele, I. & Palsson, B. Ø. What is flux balance analysis? Nat Biotechnol 28, 245–248 (2010).

7. Schellenberger, J. et al. Quantitative prediction of cellular metabolism with constraint-based models: the COBRA Toolbox v2.0. Nat Protoc 6, 1290–1307 (2011).

8. Heirendt, L. et al. Creation and analysis of biochemical constraint-based models using the COBRA Toolbox v.3.0. Nat Protoc 14, 639–702 (2019).

9. Saa, P. A. & Nielsen, L. K. Formulation, construction and analysis of kinetic models of metabolism: A review of modelling frameworks. Biotechnol Adv 35, 981–1003 (2017).

10. Strutz, J., Martin, J., Greene, J., Broadbelt, L. & Tyo, K. Metabolic kinetic modeling provides insight into complex biological questions, but hurdles remain. Curr Opin Biotech 59, 24–30 (2019).

11. Foster, C. J., Wang, L., Dinh, H. V., Suthers, P. F. & Maranas, C. D. Building kinetic models for metabolic engineering. Curr Opin Biotech 67, 35–41 (2020).

12. Srinivasan, S., Cluett, W. R. & Mahadevan, R. Constructing kinetic models of metabolism at genome-scales: A review. Biotechnol J 10, 1345–1359 (2015).

13. Liebermeister, W. & Klipp, E. Bringing metabolic networks to life: convenience rate law and thermodynamic constraints. Theor Biol Med Model 3, 41 (2006).

14. Hofmeyr, J.-H. S. & Cornish-Bowden, H. The reversible Hill equation: how to incorporate cooperative enzymes into metabolic models. Bioinformatics 13, 377–385 (1997).

15. Miskovic, L. & Hatzimanikatis, V. Modeling of uncertainties in biochemical reactions. Biotechnol Bioeng 108, 413–23 (2011).

16. Murabito, E. et al. Monte-Carlo Modeling of the Central Carbon Metabolism of Lactococcus lactis: Insights into Metabolic Regulation. Plos One 9, e106453 (2014).

17. Lee, Y., Rivera, J. G. L. & Liao, J. C. Ensemble Modeling for Robustness Analysis in engineering non-native metabolic pathways. Metab Eng 25, 63–71 (2014).

18. Khodayari, A. & Maranas, C. D. A genome-scale Escherichia coli kinetic metabolic model k-ecoli457 satisfying flux data for multiple mutant strains. Nat Commun 7, 13806 (2016).

19. Suthers, P. F., Foster, C. J., Sarkar, D., Wang, L. & Maranas, C. D. Recent advances in constraint and machine learning-based metabolic modeling by leveraging stoichiometric balances, thermodynamic feasibility and kinetic law formalisms. Metab Eng 63, 13–33 (2021).

20. Chakrabarti, A., Miskovic, L., Soh, K. C. & Hatzimanikatis, V. Towards kinetic modeling of genome-scale metabolic networks without sacrificing stoichiometric, thermodynamic and physiological constraints. Biotechnol J 8, 1043–1057 (2013).

21. Schmidhuber, J. Deep learning in neural networks: An overview. Neural Networks 61, 85–117 (2015).

22. Goodfellow, I. J. et al. Generative Adversarial Networks. Arxiv (2014).

23. Moret, M., Friedrich, L., Grisoni, F., Merk, D. & Schneider, G. Generative molecular design in low data regimes. Nat Mach Intell 2, 171–180 (2020).

24. Pan, S. J. & Yang, Q. A Survey on Transfer Learning. Ieee T Knowl Data En 22, 1345–1359 (2009).

25. Miskovic, L. & Hatzimanikatis, V. Production of biofuels and biochemicals: in need of an ORACLE. Trends Biotechnol 28, 391–397 (2010).

26. Mirza, M. & Osindero, S. Conditional Generative Adversarial Nets. Arxiv (2014).

27. Henry, C. S., Broadbelt, L. J. & Hatzimanikatis, V. Thermodynamics-based metabolic flux analysis. Biophys J 92, 1792–805 (2006).

28. Ataman, M. & Hatzimanikatis, V. Heading in the right direction: thermodynamics-based network analysis and pathway engineering. Curr Opin Biotech 36, 176–182 (2015).

29. Salvy, P. et al. pyTFA and matTFA: a Python package and a Matlab toolbox for Thermodynamics-based Flux Analysis. Bioinformatics 35, 167–169 (2019).

30. Varma, A. & Palsson, B. O. Metabolic Capabilities of Escherichia coli: I. Synthesis of Biosynthetic Precursors and Cofactors. J Theor Biol 165, 477–502 (1993).

31. Soh, K. C., Miskovic, L. & Hatzimanikatis, V. From network models to network responses: integration of thermodynamic and kinetic properties of yeast genome-scale metabolic networks. Fems Yeast Res 12, 129–143 (2012).

32. Andreozzi, S. et al. Identification of metabolic engineering targets for the enhancement of 1,4-butanediol production in recombinant E. coli using large-scale kinetic models. Metab Eng 35, 148–159 (2016).

33. Miskovic, L. et al. A design–build–test cycle using modeling and experiments reveals interdependencies between upper glycolysis and xylose uptake in recombinant S. cerevisiae and improves predictive capabilities of large-scale kinetic models. Biotechnol Biofuels 10, 166 (2017).

34. Hameri, T., Fengos, G., Ataman, M., Miskovic, L. & Hatzimanikatis, V. Kinetic models of metabolism that consider alternative steady-state solutions of intracellular fluxes and concentrations. Metab Eng 52, 29–41 (2019).

35. Tokic, M., Hatzimanikatis, V. & Miskovic, L. Large-scale kinetic metabolic models of Pseudomonas putida KT2440 for consistent design of metabolic engineering strategies. Biotechnol Biofuels 13, 33 (2020).

36. Weilandt, D. & al., et. Symbolic Kinetic Models in Python (SKiMpy): Intuitive modeling of large scale biological kinetic models. in preparation (2021).

37. Srivastava, A., Valkov, L., Russell, C., Gutmann, M. U. & Sutton, C. VEEGAN: Reducing Mode Collapse in GANs using Implicit Variational Learning. Arxiv (2017).

38. Miskovic, L., Béal, J., Moret, M. & Hatzimanikatis, V. Uncertainty reduction in biochemical kinetic models: Enforcing desired model properties. Plos Comput Biol 15, e1007242 (2019).

39. Andreozzi, S., Miskovic, L. & Hatzimanikatis, V. iSCHRUNK – In Silico Approach to Characterization and Reduction of Uncertainty in the Kinetic Models of Genome-scale Metabolic Networks. Metab Eng 33, 158–168 (2016).

40. Gutenkunst, R. N. et al. Universally Sloppy Parameter Sensitivities in Systems Biology Models. Plos Comput Biol 3, e189 (2007).

41. McInnes, L., Healy, J., Saul, N. & Großberger, L. UMAP: Uniform Manifold Approximation and Projection. J Open Source Softw 3, 861 (2018).

42. Weiss, K., Khoshgoftaar, T. M. & Wang, D. A survey of transfer learning. J Big Data 3, 9 (2016).

43. Che, T., Li, Y., Jacob, A. P., Bengio, Y. & Li, W. Mode Regularized Generative Adversarial Networks. Arxiv (2016).

44. Torrie, G. M. & Valleau, J. P. Nonphysical sampling distributions in Monte Carlo free-energy estimation: Umbrella sampling. J Comput Phys 23, 187–199 (1977).

45. Kroese, D. P., Brereton, T., Taimre, T. & Botev, Z. I. Why the Monte Carlo method is so important today. Wiley Interdiscip Rev Comput Statistics 6, 386–392 (2014).

46. Miskovic, L., Tokic, M., Savoglidis, G. & Hatzimanikatis, V. Control Theory Concepts for Modeling Uncertainty in Enzyme Kinetics of Biochemical Networks. Ind Eng Chem Res 58, 13544–13554 (2019).

47. Wang, L., Birol, I. & Hatzimanikatis, V. Metabolic Control Analysis under Uncertainty: Framework Development and Case Studies. Biophys J 87, 3750–3763 (2004).

48. Soh, K. C. & Hatzimanikatis, V. Network thermodynamics in the post-genomic era. Curr Opin Microbiol 13, 350–357 (2010).

49. Soh, K. C. & Hatzimanikatis, V. Constraining the flux space using thermodynamics and integration of metabolomics data. Methods Mol Biology Clifton N J 1191, 49–63 (2014).

50. Dunteman, G. Principal Components Analysis. (1989) doi:10.4135/9781412985475.

51. Hinton, L. van der M. and G. E. Visualizing Data using t-SNE. JMLR 9, 2579–2605 (2008).

52. Kullback, S. & Leibler, R. A. On Information and Sufficiency. Ann Math Statistics 22, 79–86 (1951).

53. Abadi, M. et al. TensorFlow: A system for large-scale machine learning. Arxiv (2016).

54. Kingma, D. P. & Ba, J. Adam: A Method for Stochastic Optimization. Arxiv (2014).

