## Supplementary information for "Reconstructing Kinetic Models for Dynamical Studies of Metabolism using Generative Adversarial Networks"

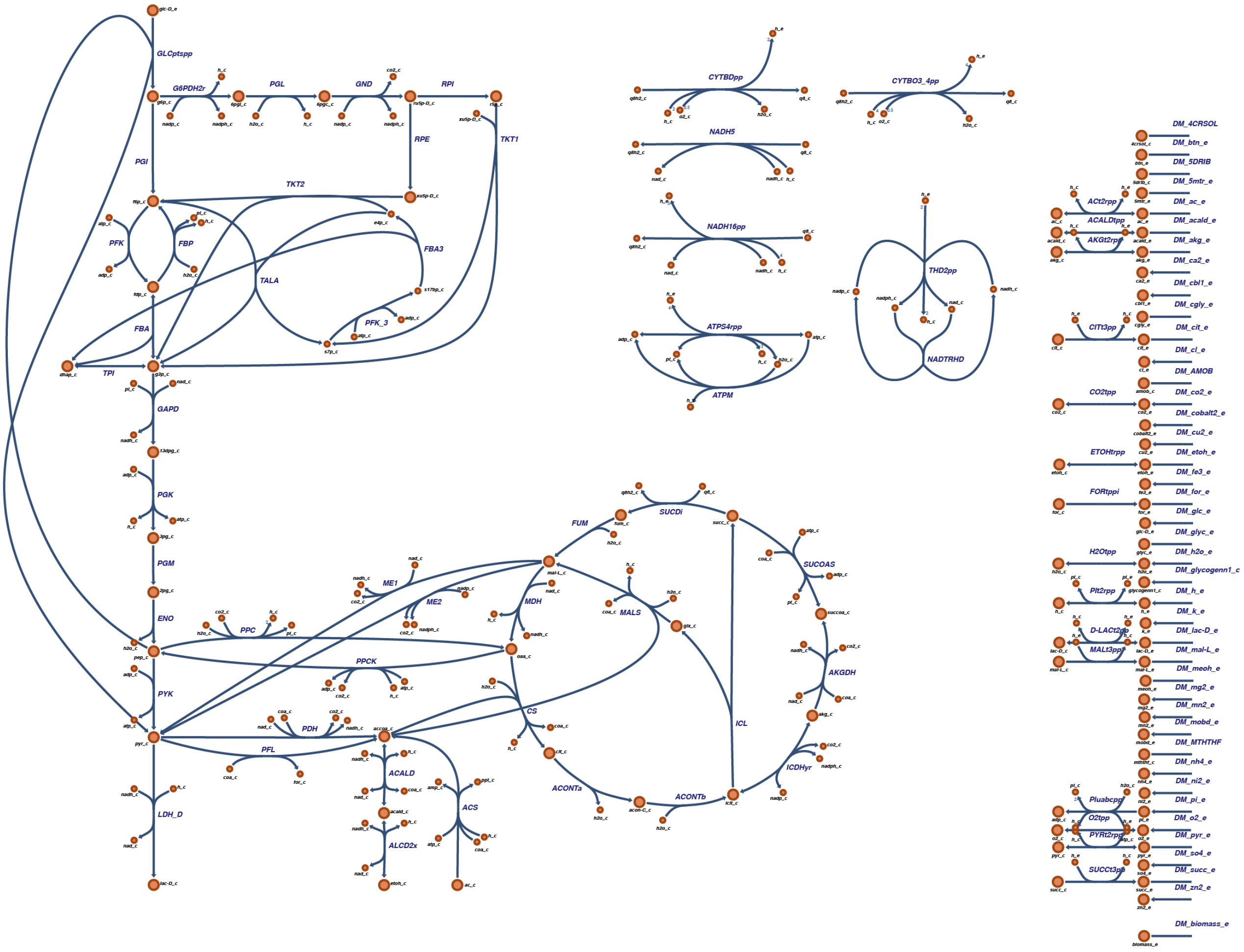


**Supplementary Figure 1:** Escher map of the *E. coli* central carbon metabolism network. The reactions are represented as blue lines and the metabolites as orange circles. This map was generated using the Escher web tool[1].

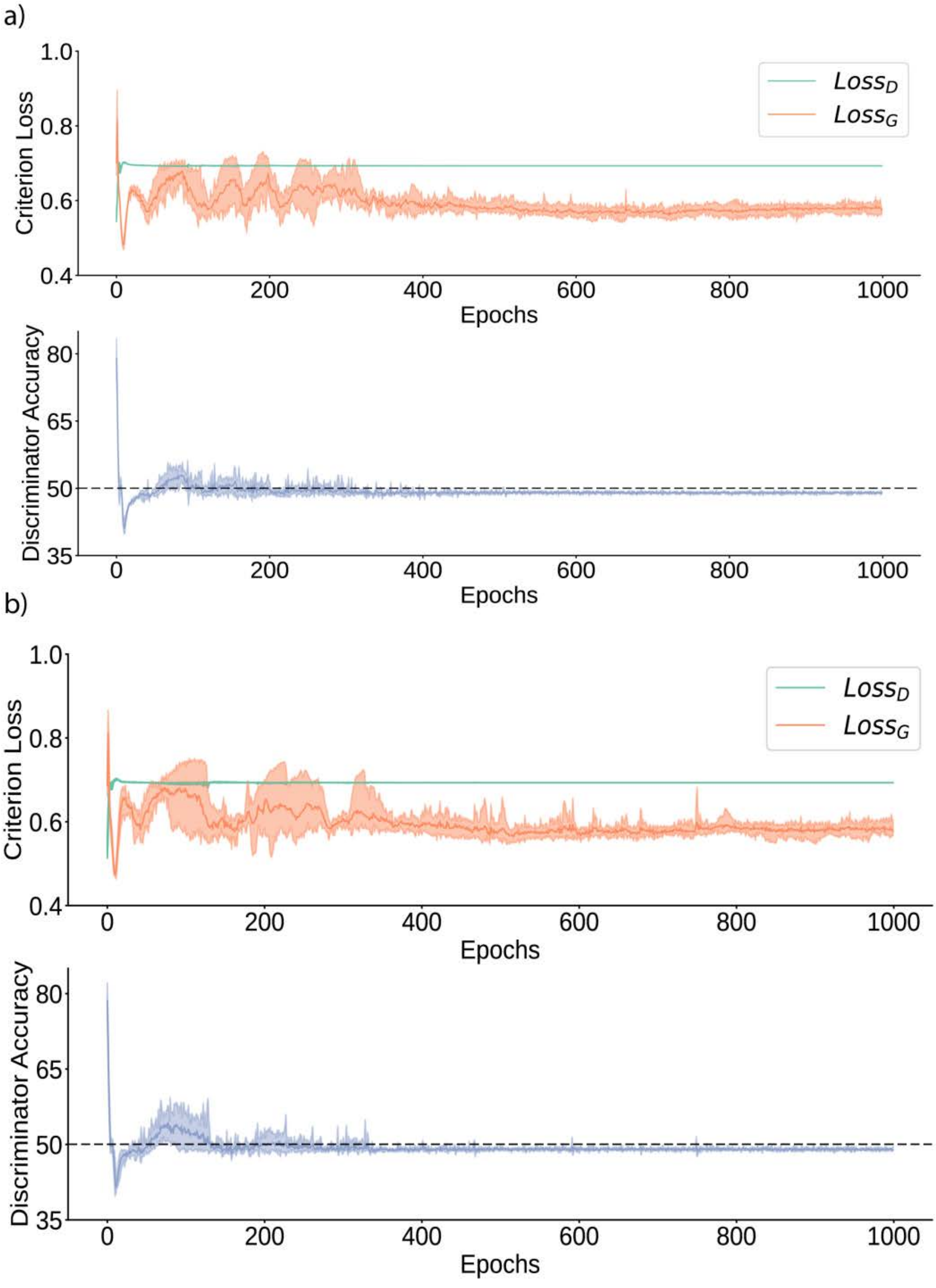
a)
b)
Criterion Loss
Discriminator Accuracy
Epochs
$Loss_D$
$Loss_G$
1.0
0.8
0.6
0.4
80
65
50
35
0
200
400
600
800
1000

c)

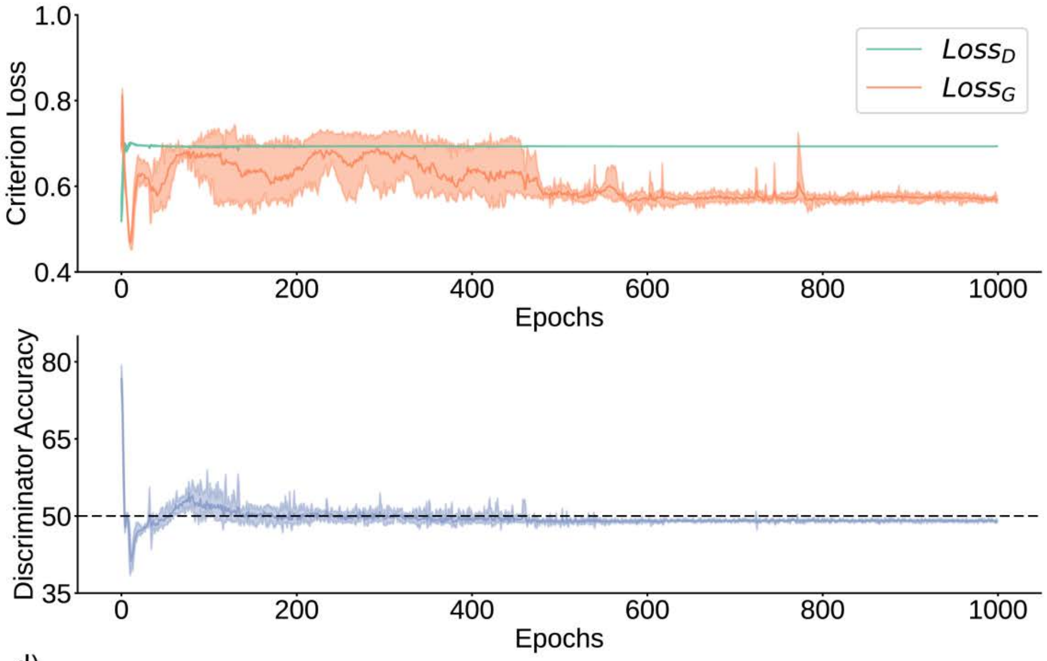


d)

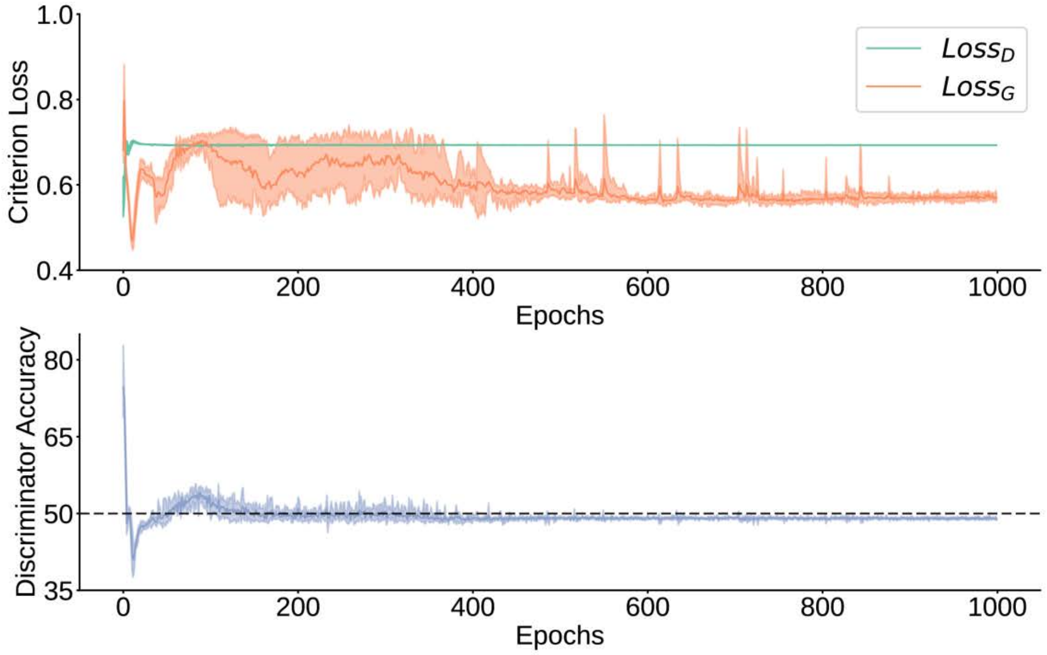


**Supplementary Figure 2:** Upper Panel: Discriminator Loss (green) and generator Loss (orange) values over training epochs for 5 statistical repeats. Lower Panel: Discriminator Accuracy over training epochs (5 statistical repeats) for (a) Physiology 1 (b) Physiology 2 (c) Physiology 3 (d) Physiology 4. The losses of the discriminator and generator stabilize as training progresses. The discriminator accuracy settles near 50% indicating that the discriminator cannot distinguish between training data (from ORACLE) and generated data (from the generator).

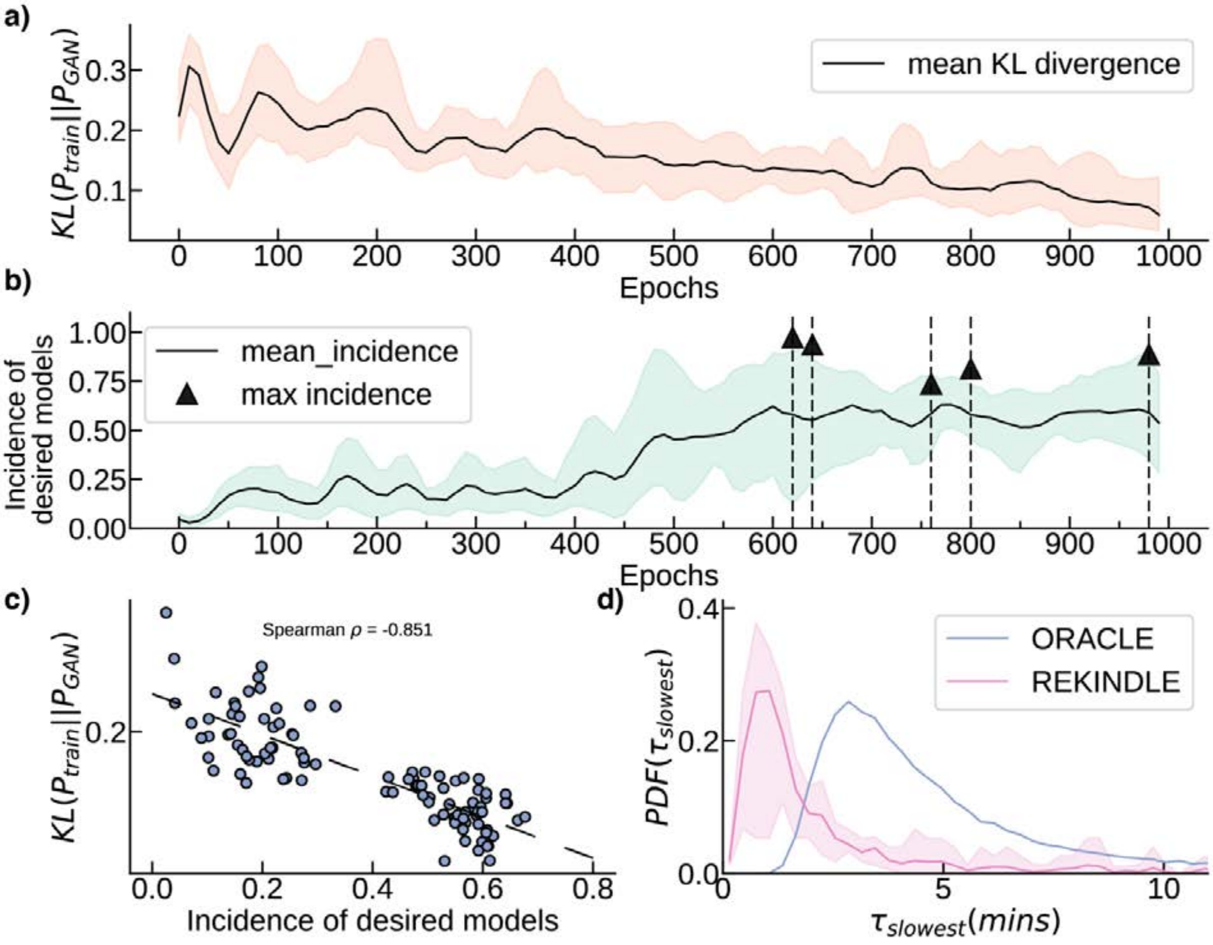


**Supplementary Figure 3a:** Generation and Validation of REKINDLE generated datasets for Physiology 2. The panels description and the meaning of abbreviations are presented in the caption of Figure 2 of the main manuscript.

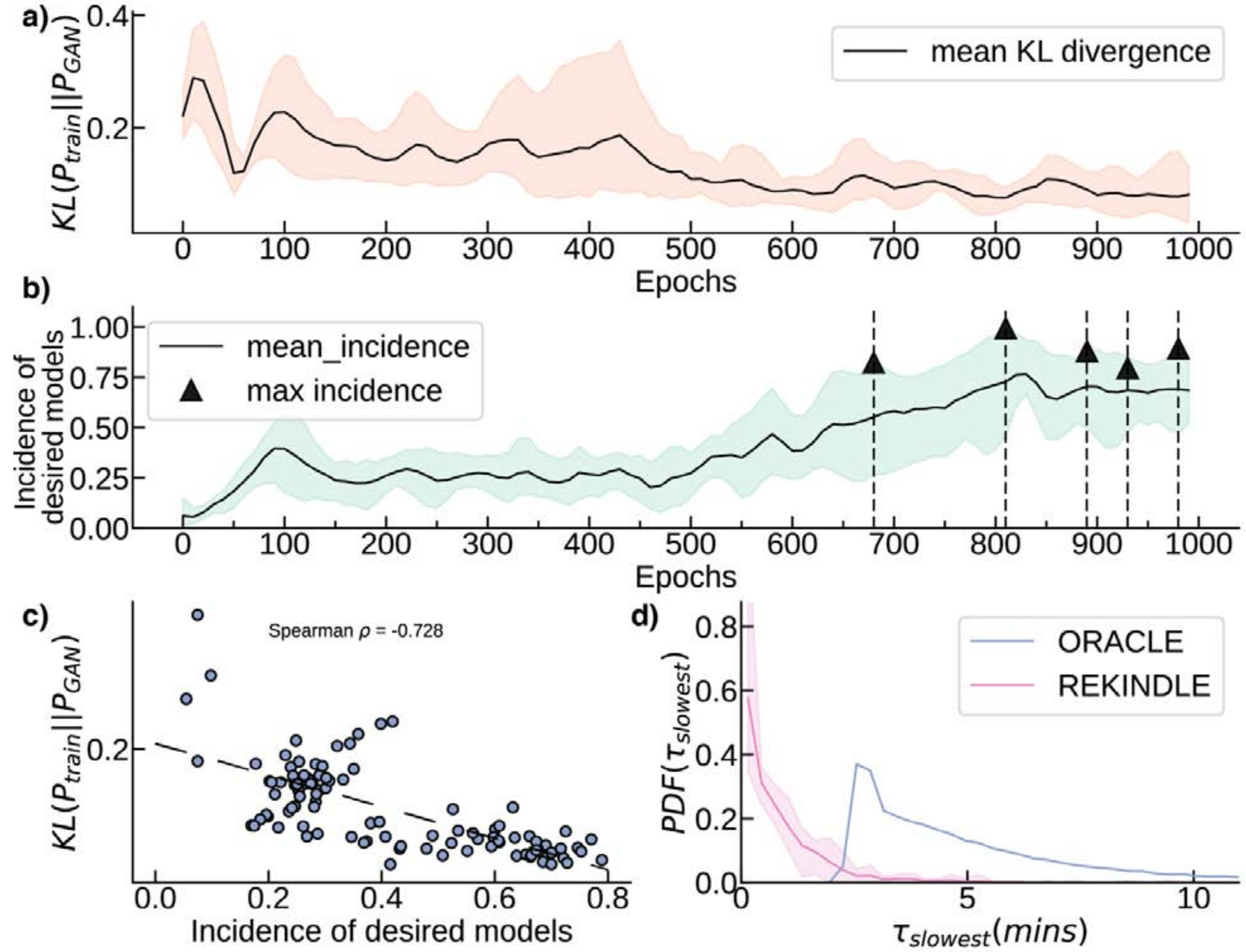


**Supplementary Figure 3b:** Generation and Validation of REKINDLE generated datasets for Physiology 3. The panels description and the meaning of abbreviations are presented in the caption of Figure 2 of the main manuscript.

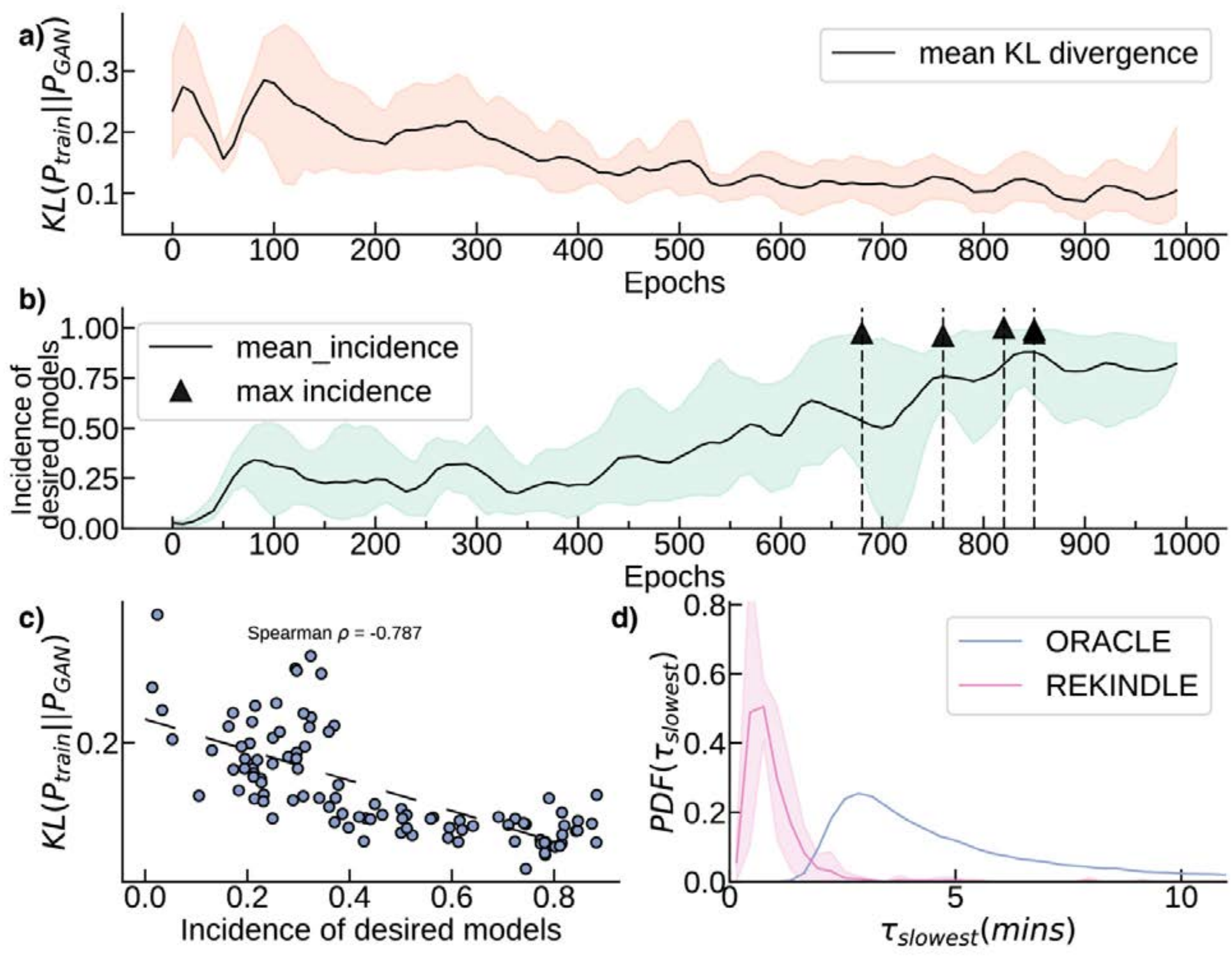


**Supplementary Figure 3c:** Generation and Validation of REKINDLE generated datasets for Physiology 4. The panels description and the meaning of abbreviations are presented in the caption of Figure 2 of the main manuscript.

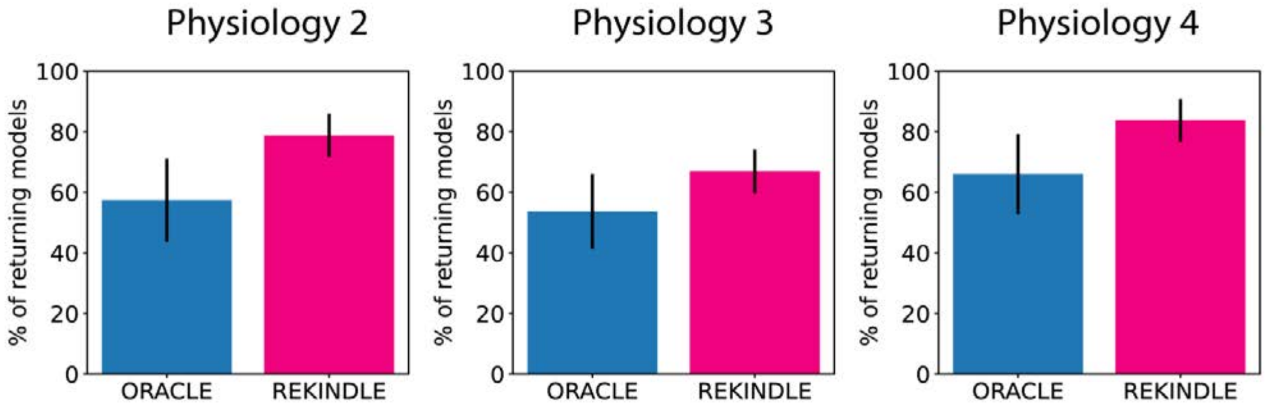


**Supplementary Figure 4:** 1000 models were perturbed 10 times randomly between $0.5X_{RSS} \leq \Delta X \leq 2X_{RSS}$ and then allowed to evolve with the perturbed state, $\Delta X$, as the initial condition and then checked if the dynamics comes back to within 1% of the reference steady state for (a) Physiology 2 (b) Physiology 3 (c) Physiology 4. We see that higher percentage of REKINDLE parameterized kinetic models comes back to the steady state compared to ORACLE.

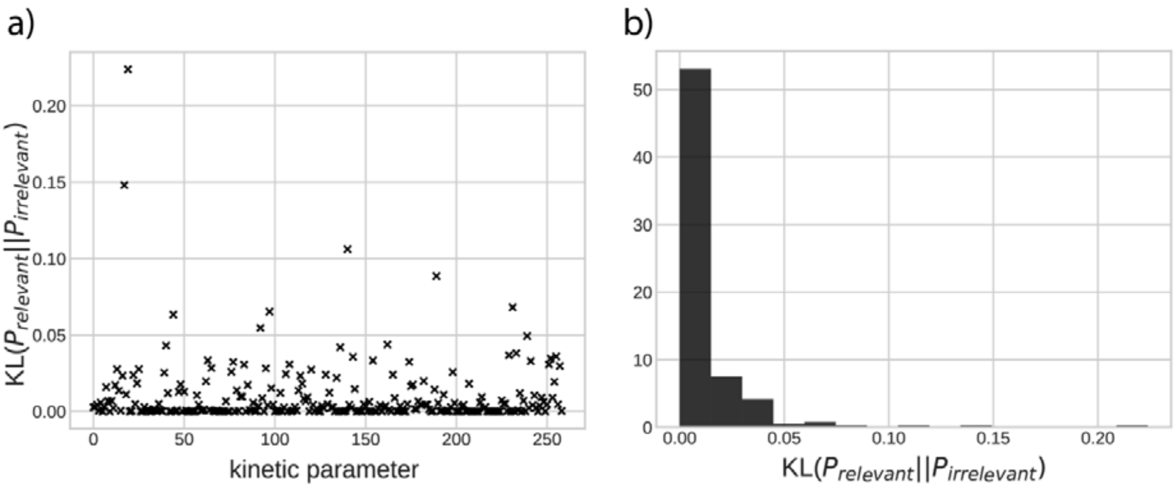


**Supplementary Figure 5:** (a) KL divergence scores of kinetic parameters distributions with biologically relevant and non relevant dynamics for Physiology 1. (b) Distributions of the KL divergence scores in (a). We observe that only a handful of parameters have significant KL divergence scores and majority of the parameters have KL divergence score close to zero. This indicates the most parameters have the same distributions in the kinetic models with relevant and non-relevant dynamics.

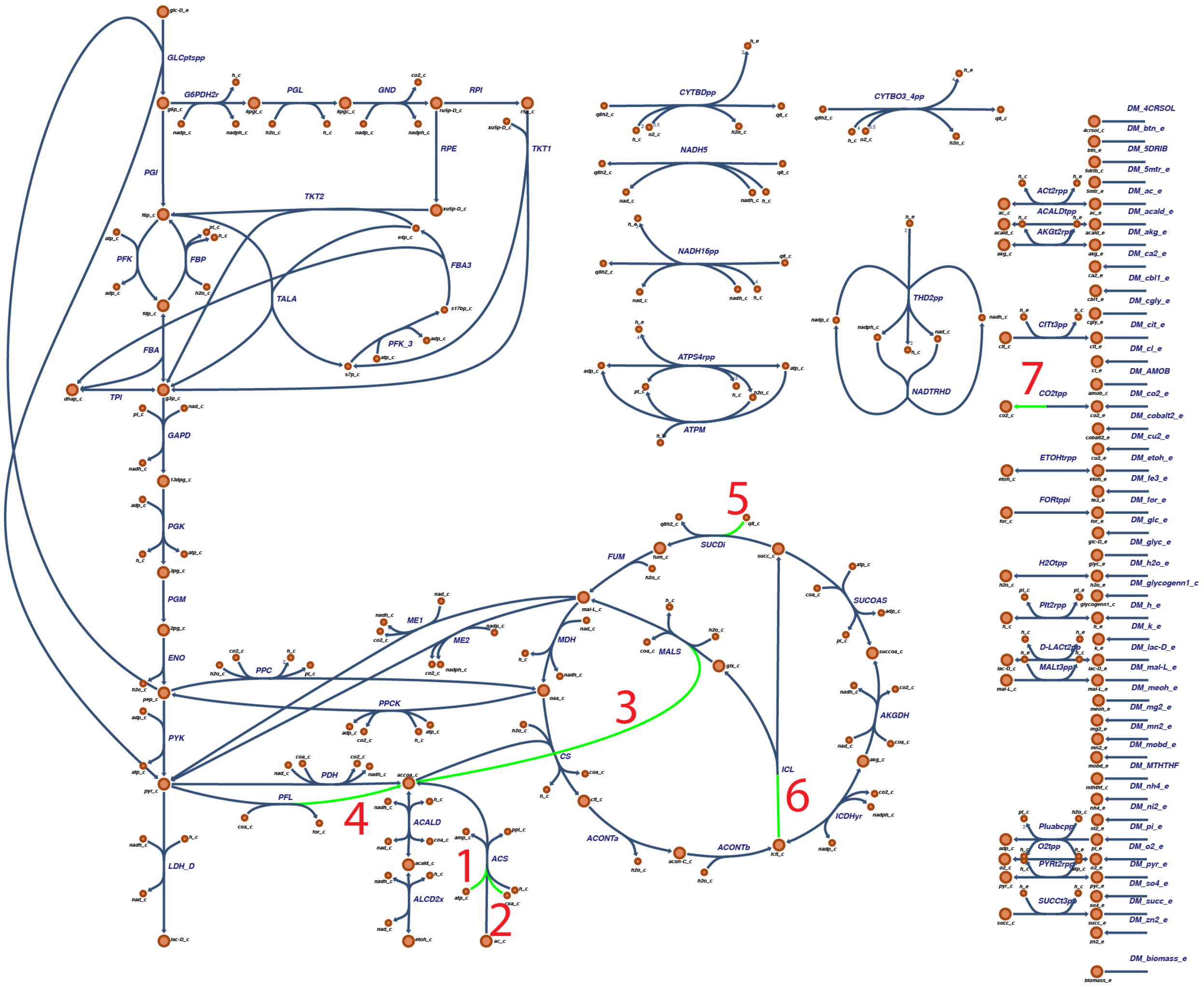


**Supplementary Figure 6:** The top 7 kinetic parameters from Fig 3a are visualized in the metabolic network. The green highlight indicates the substrate of the kinetic parameter with their associated reaction. The red numbers indicate the rank of the kinetic parameter according to Figure 3a. The exact reactions and rate law expressions of these 7 parameters are elaborated in Supplementary Table S1.

| Kinetic Parameter | Reaction Name, Reaction | Rate law repression |
|---|---|---|
| Rank 1, $K_{M,atp}^{ACS}$ | ACS: $coa_c+ac_c+atp_c \rightleftharpoons accoa_c + ppi_c+amp_c$ | $v = \frac{V_{max}\left(1-\frac{1}{K_{eq}}\prod_{i=1}^{N}\left(\frac{[P_i]}{[S_i]}\right)\right)\prod_{i=1}^{N}\left(\frac{[S_i]}{K_{M,S_i}}\right)\left(\frac{[S_i]}{K_{M,S_i}}+\frac{[P_i]}{K_{M,P_i}}\right)^{h-1}}{\prod_{i=1}^{N}\left(1+\left(\frac{[S_i]}{K_{M,S_i}}+\frac{[P_i]}{K_{M,P_i}}\right)^{h}\right)}$<br>Parameters:<br>$[V_{max}, K_{M,coa}^{ACS}, K_{M,ac}^{ACS}, K_{M,atp}^{ACS}, K_{M,accoa}^{ACS}, K_{M,ppi}^{ACS}, K_{M,amp}^{ACS}, K_{eq}, h, N = 3]$ |
| Rank 2, $K_{M,coa}^{ACS}$ | ACS: $coa_c+ac_c+atp_c \rightleftharpoons accoa_c + ppi_c+amp_c$ | $v = \frac{V_{max}\left(1-\frac{1}{K_{eq}}\prod_{i=1}^{N}\left(\frac{[P_i]}{[S_i]}\right)\right)\prod_{i=1}^{N}\left(\frac{[S_i]}{K_{M,S_i}}\right)\left(\frac{[S_i]}{K_{M,S_i}}+\frac{[P_i]}{K_{M,P_i}}\right)^{h-1}}{\prod_{i=1}^{N}\left(1+\left(\frac{[S_i]}{K_{M,S_i}}+\frac{[P_i]}{K_{M,P_i}}\right)^{h}\right)}$<br>Parameters:<br>$[V_{max}, K_{M,coa}^{ACS}, K_{M,ac}^{ACS}, K_{M,atp}^{ACS}, K_{M,accoa}^{ACS}, K_{M,ppi}^{ACS}, K_{M,amp}^{ACS}, K_{eq}, h, N = 3]$ |
| Rank 3, $K_{M,accoa}^{MALS}$ | MALS: $glx_c+accoa_c \rightleftharpoons mal_{L_c} + coa_c$ | $v = \frac{V_{max}\left(1-\frac{1}{K_{eq}}\prod_{i=1}^{N}\left(\frac{[P_i]}{[S_i]}\right)\right)\prod_{i=1}^{N}\left(\frac{[S_i]}{K_{M,S_i}}\right)\left(\frac{[S_i]}{K_{M,S_i}}+\frac{[P_i]}{K_{M,P_i}}\right)^{h-1}}{\prod_{i=1}^{N}\left(1+\left(\frac{[S_i]}{K_{M,S_i}}+\frac{[P_i]}{K_{M,P_i}}\right)^{h}\right)}$<br>Parameters: $[V_{max}, K_{M,glx}^{MALS}, K_{M,accoa}^{MALS}, K_{M,mal_{L_c}}^{MALS}, K_{M,coa}^{MALS},, K_{eq}, h, N = 2]$ |

| | | |
|---|---|---|
| Rank 4, $K_{M,accoa}^{PFL}$ | PFL: $coa_c + pyr_c \rightleftharpoons for_c + accoa_c$ | $v = \frac{V_{max}\left(1-\frac{1}{K_{eq}}\prod_{i=1}^{N}\left(\frac{[P_i]}{[S_i]}\right)\right)\prod_{i=1}^{N}\left(\frac{[S_i]}{K_{M,S_i}}\right)\left(\frac{[S_i]}{K_{M,S_i}}+\frac{[P_i]}{K_{M,P_i}}\right)^{h-1}}{\prod_{i=1}^{N}\left(1+\left(\frac{[S_i]}{K_{M,S_i}}+\frac{[P_i]}{K_{M,P_i}}\right)^{h}\right)}$<br>Parameters: $[V_{max}, K_{M,coa}^{PFL}, K_{M,pyr}^{PFL}, K_{M,for_c}^{PFL}, K_{M,accoa}^{PFL}, K_{eq}, h, N = 2]$ |
| Rank 5, $K_{M,q8}^{SUCDi}$ | SUCDi: $q8_c + succ_c \rightleftharpoons fum_c + q8h2_c$ | $v = \frac{V_{max}\left(1-\frac{1}{K_{eq}}\prod_{i=1}^{N}\left(\frac{[P_i]}{[S_i]}\right)\right)\prod_{i=1}^{N}\left(\frac{[S_i]}{K_{M,S_i}}\right)\left(\frac{[S_i]}{K_{M,S_i}}+\frac{[P_i]}{K_{M,P_i}}\right)^{h-1}}{\prod_{i=1}^{N}\left(1+\left(\frac{[S_i]}{K_{M,S_i}}+\frac{[P_i]}{K_{M,P_i}}\right)^{h}\right)}$<br>Parameters: $[V_{max}, K_{M,q8}^{SUCDi}, K_{M,succ}^{SUCDi}, K_{M,fum}^{SUCDi}, K_{M,q8h2}^{SUCDi}, K_{eq}, h, N = 2]$ |
| Rank 6, $K_{M,icit}^{ICL}$ | ICL: $icit_c \rightleftharpoons succ_c + glx_c$ | $v = \frac{V_{max}\prod_{i=1}^{M}\left(\frac{[S_i]}{K_{M,S_i}}\right)\left(1-\frac{1}{K_{eq}}\frac{\prod_{j=1}^{N}[P_j]}{\prod_{i=1}^{M}[S_i]}\right)}{\prod_{i=1}^{M}\sum_{m=0}^{\alpha_i}\left(\frac{[S_i]}{K_{M,S_i}}\right)^{m}+\prod_{j=1}^{N}\sum_{m=0}^{\beta_j}\left(\frac{[P_j]}{K_{M,P_j}}\right)^{m}-1}$<br>Parameters: $[V_{max}, K_{M,icit}^{ICL}, K_{M,succ}^{ICL}, K_{M,q8h2}^{ICL}, K_{eq}, N = 2, M = 1, \alpha_1 = 1, \beta_1 = -1, \beta_2 = -1]$ |
| Rank 7, $K_{M,co2}^{CO2tpp}$ | CO2tpp: $co2_e \rightleftharpoons co2_c$ | $v = \frac{V_{max}\left(1-\frac{1}{K_{eq}}\prod_{i=1}^{N}\left(\frac{[P_i]}{[S_i]}\right)\right)\prod_{i=1}^{N}\left(\frac{[S_i]}{K_{M,S_i}}\right)\left(\frac{[S_i]}{K_{M,S_i}}+\frac{[P_i]}{K_{M,P_i}}\right)^{h-1}}{\prod_{i=1}^{N}\left(1+\left(\frac{[S_i]}{K_{M,S_i}}+\frac{[P_i]}{K_{M,P_i}}\right)^{h}\right)}$<br>Parameters: $[V_{max}, K_{M,co2_c}^{CO2tpp}, K_{M,co2_e}^{CO2tpp}\ K_{eq}, h, N = 1]$ |

**Supplementary Table 1:** The reaction and the rate law formalisms of the top 7 kinetic parameters from Fig. 3a. Here $S_i$ and $P_i$ represents the $i^{th}$ substrate and product of a reaction, $K_{eq}$ represents the equilibrium constant of the reaction and $h$ is the hill coefficient of the generalized Michaelis-Menten and Convenience kinetics .

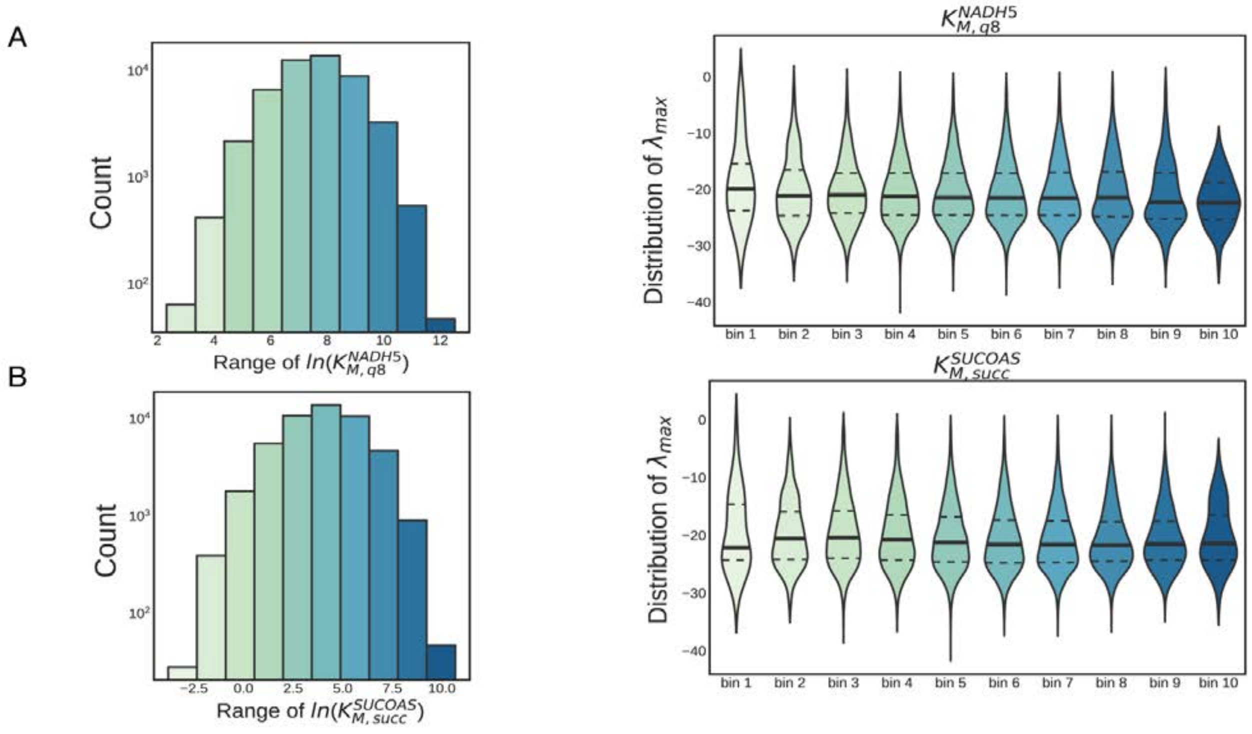


**Supplementary Figure 7:** 50,000 REKINDLE generated models were divided into 10 sub-populations (indicated by the different shades of the bins) based on their respective values of A) $K_{M,q8}^{NADH5}$ B) $K_{M,suc}^{SUCOAS}$ (left). and (Right) the maximum eigenvalue distributions of the Jacobian for the sub-populations. The thick black line indicates the mean of the distribution and the dashed lines indicates quartiles. These are the lowest ranked parameters in Fig 3a and consequently show negligible effect on the dynamics compared to high ranked parameters.

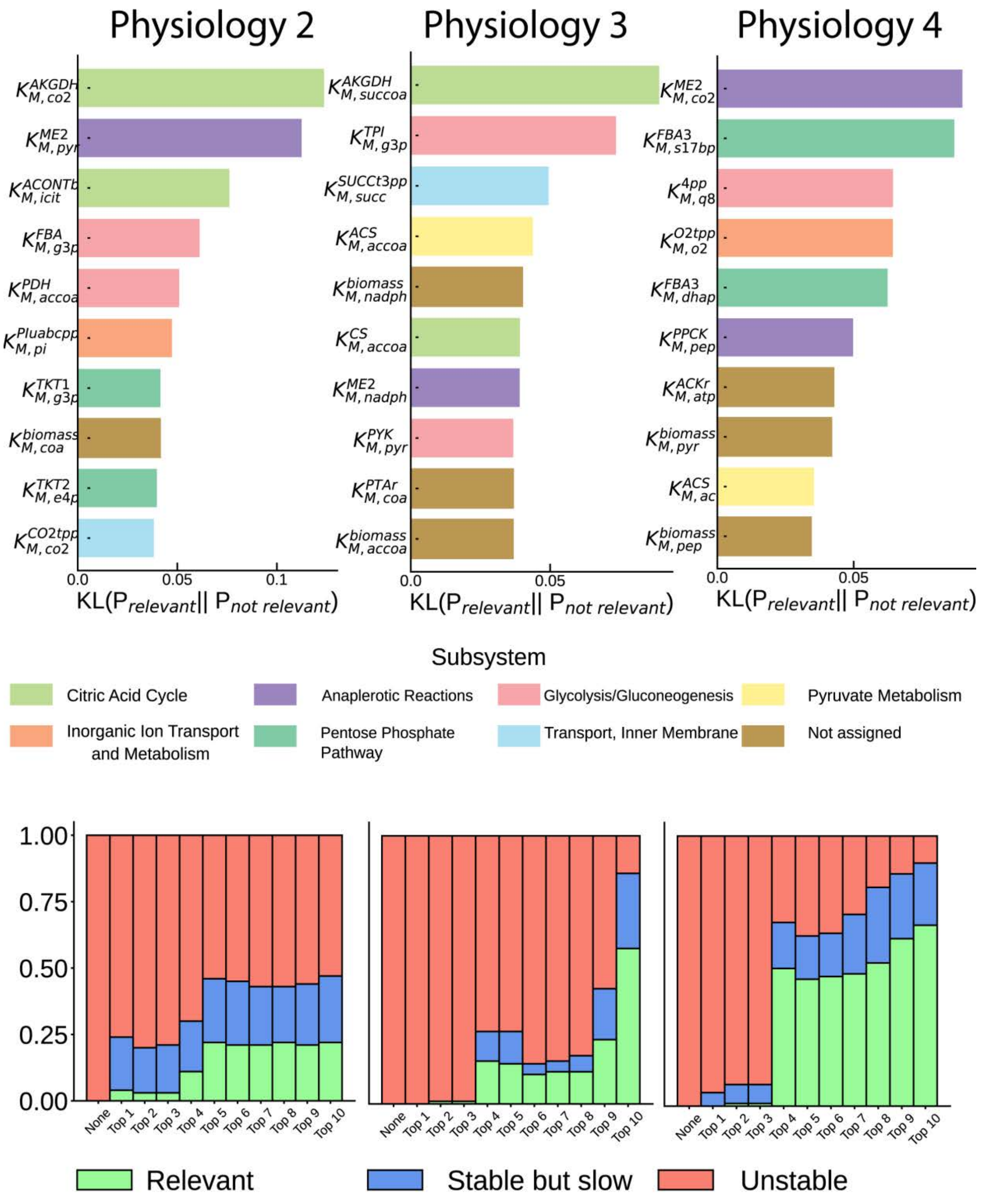


**Supplementary Figure 8:** Top: KL divergence score between distributions of the relevant and non-relevant class of generated kinetic parameters for Physiology 2 (left), Physiology 3 (middle), Physiology 4 (right) and the respective metabolic subsystems thy belong to. Bottom: We begin with 100 locally unstable models (orange bar, left) for each physiology and gradually constrain the ranked parameters in the top panel in a cumulative manner, based on their ranks (right). Constraining the parameters stabilizes (blue) as well as rescues (green) the unstable models.

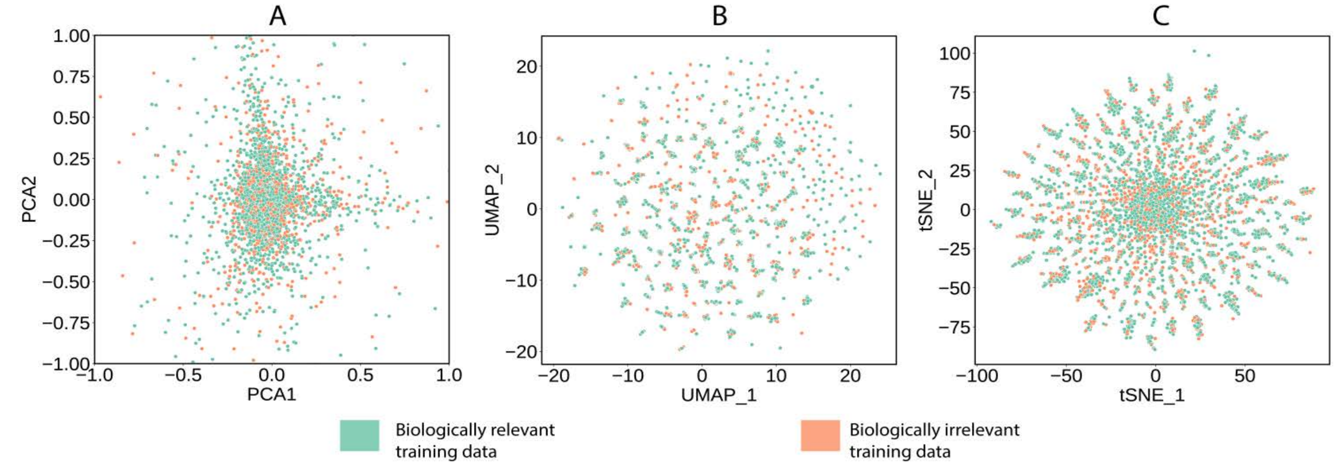


**Supplementary Figure 9:** The kinetic parameter space of the training dataset for Physiology 1 was visualized using different unsupervised dimension reduction tools (a) PCA (b) UMAP (c) tSNE. We observe that these tools are not able to recover a proper boundary between the relevant and non-relevant kinetic parameter sets. We observed similar results for all the 4 different physiologies.

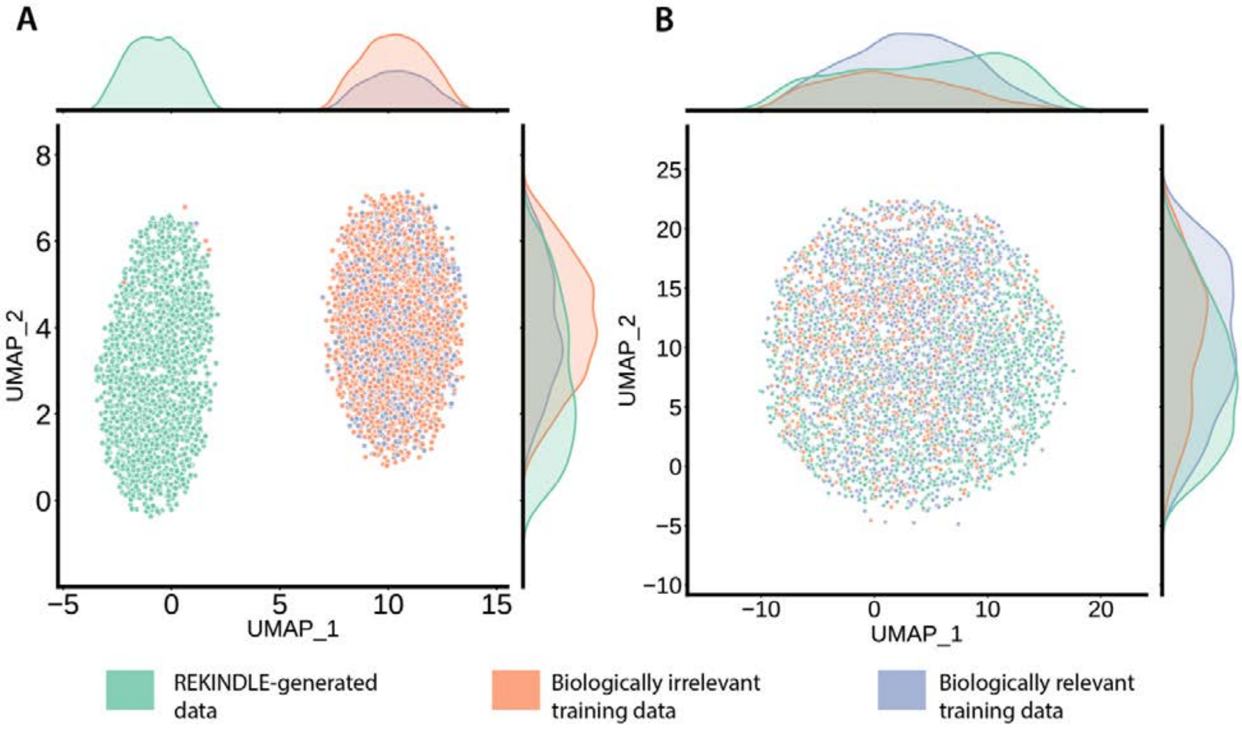


**Supplementary Figure 10:** The kinetic parameter space was visualized using UMAP (a) with all the parameters (b) Only the top 5 parameters in Fig 3a. We observe that in the (a) the REKINDLE generated data (green) and the training set (orange and blue) are completely disjoint and in (b) they share the same space. This indicates that the GANs in REKINDLE are able to distinguish between significant parameters and non-significant parameters and learn them accordingly.

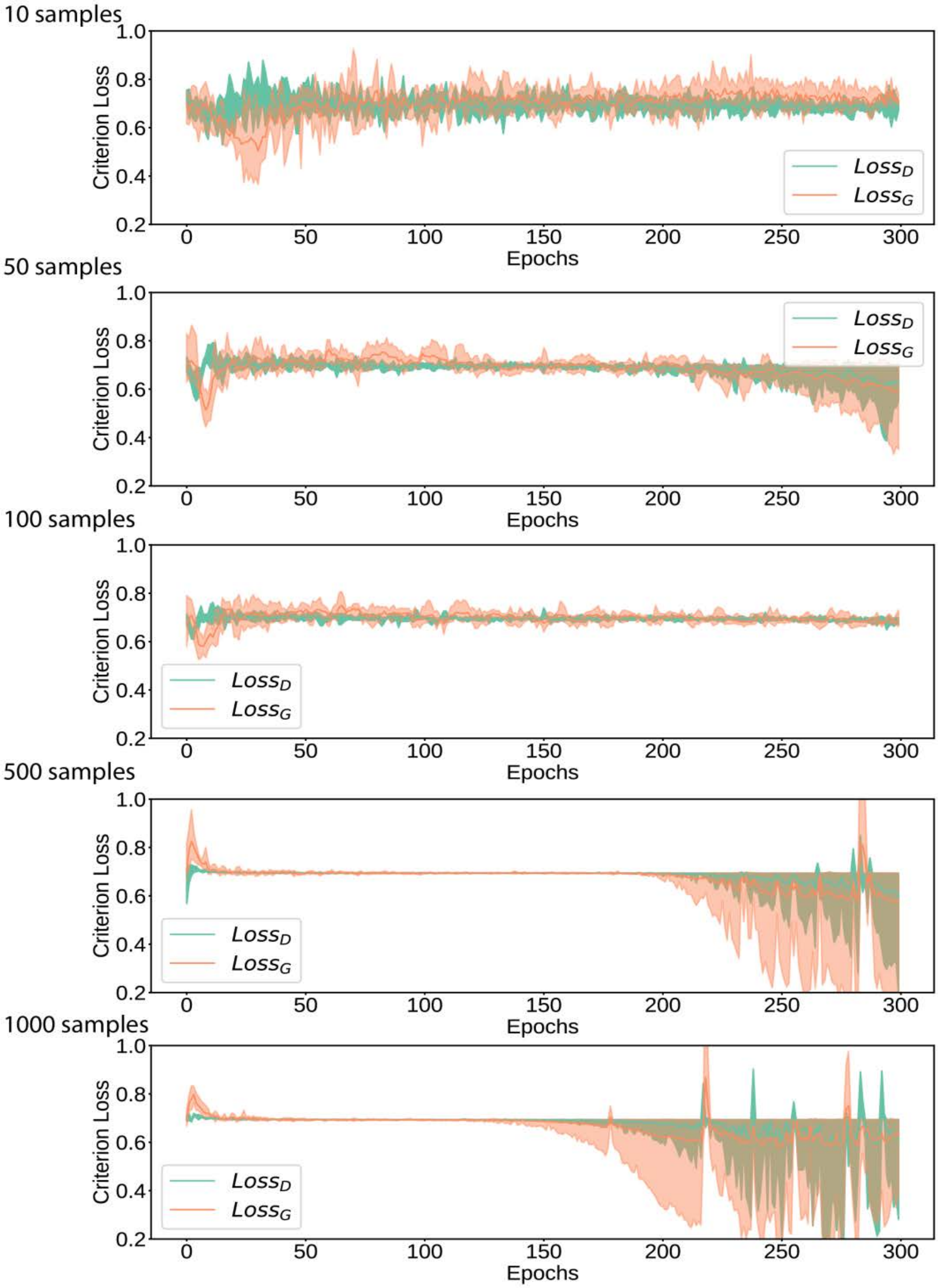


**Supplementary Figure 11a:** The discriminator loss (green line) and generator loss (orange line) for the case of transfer learning from Physiology 4 to Physiology 1, using 10 (top panel), 50, 100, 500 and 1000 (bottom panel) samples from physiology 1, for 5 statistical repeats. The losses show similar trends for all 11 other cases of transfer learning.

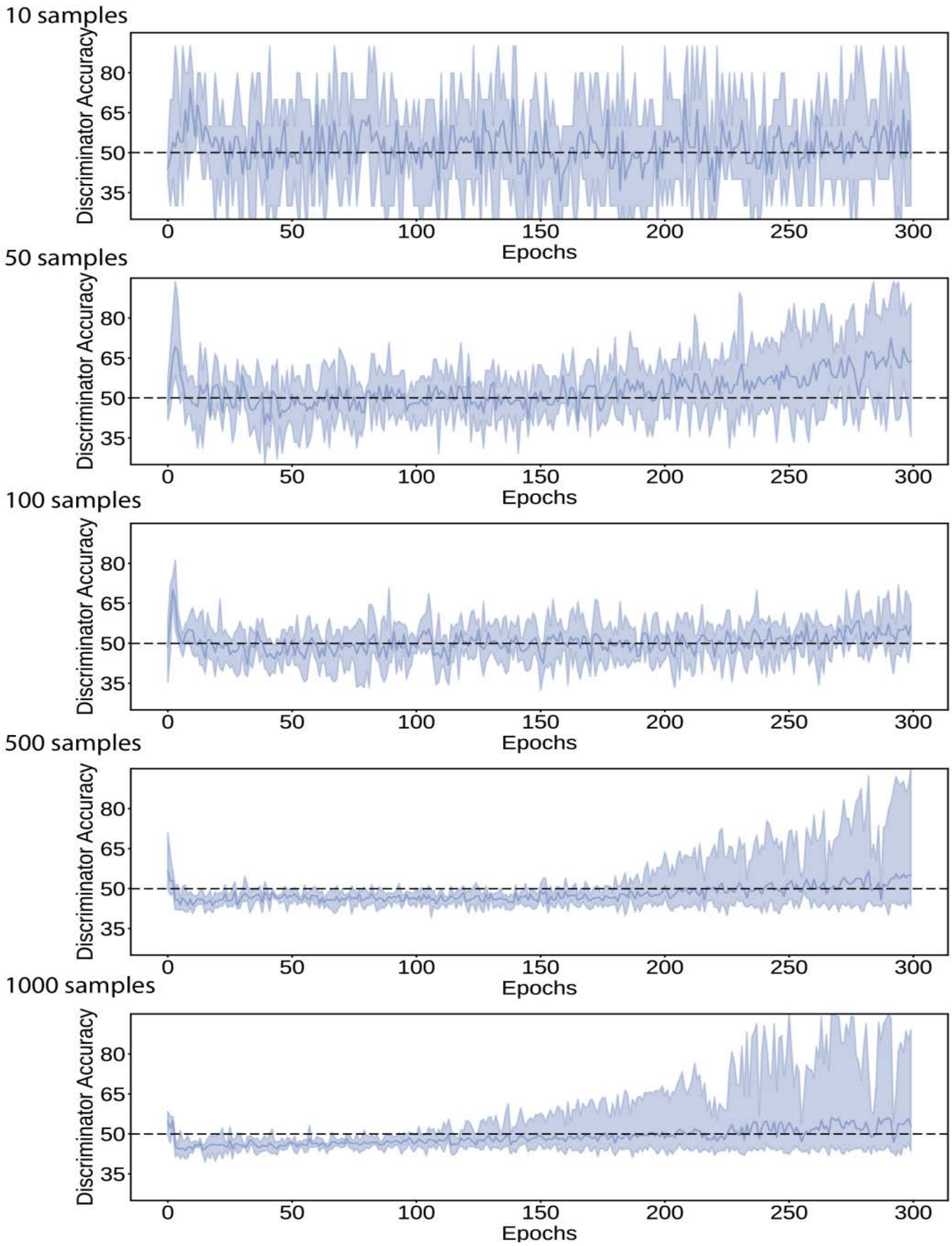


**Supplementary Figure 11b:** The discriminator accuracy for the case of transfer learning from Physiology 4 to Physiology 1, using 10 (top panel), 50, 100, 500 and 1000 (bottom panel) samples from physiology 1, for 5 statistical repeats. The accuracies show similar trends for all 11 other cases of transfer learning. It can be seen that training beyond 150 epochs leads to discriminator overpowering the generator for the cases with 500 and 1000 samples.

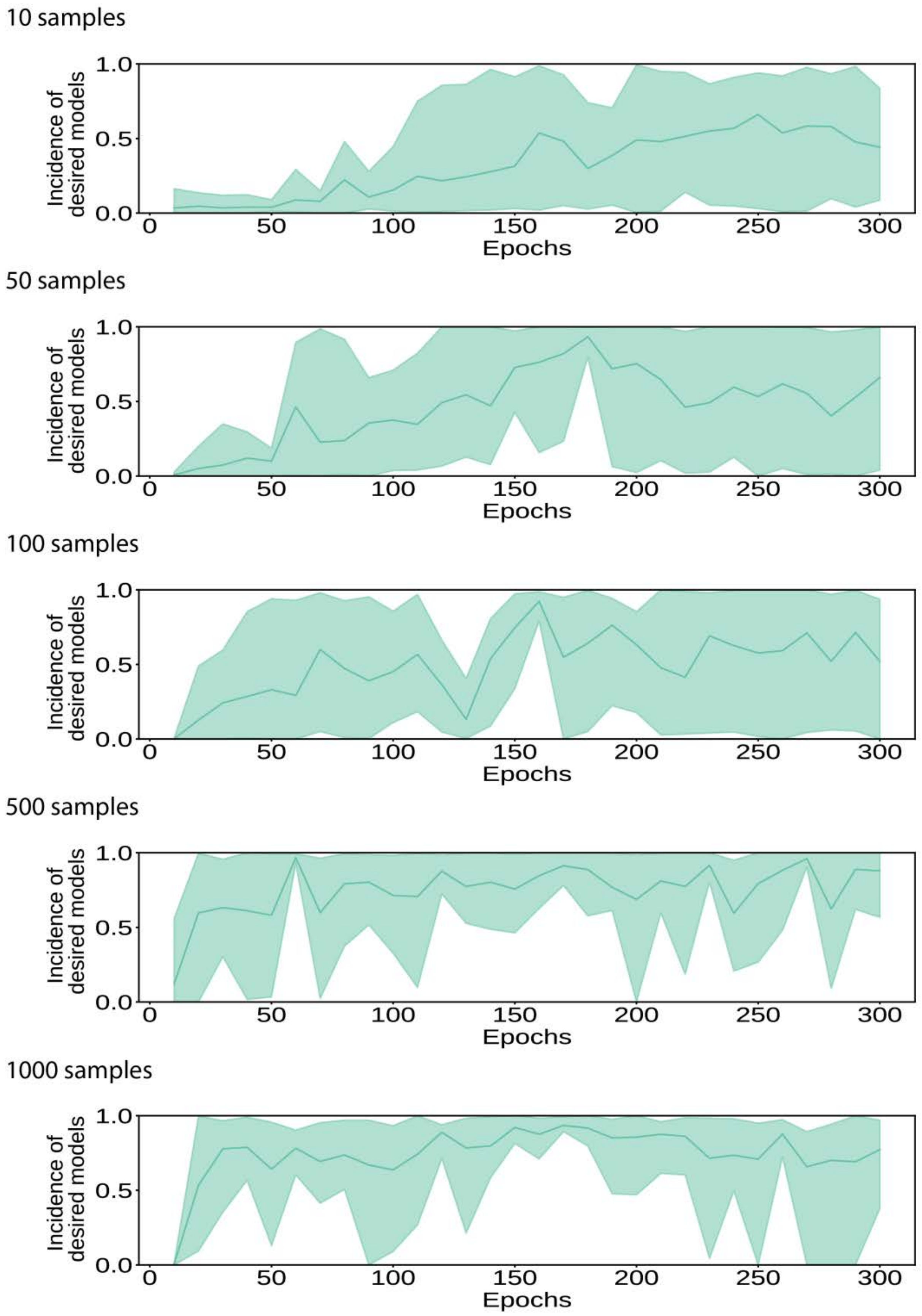


**Supplementary Figure 11c:** The incidence of biologically relevant models for the case of transfer learning from Physiology 4 to Physiology 1, using 10 (top panel), 50, 100, 500 and 1000 (bottom panel) samples from physiology 1, for 5 statistical repeats. The incidences show similar trends for all 11 other cases of transfer learning.

### Supplementary Note 1: Generating biologically non-relevant models with REKINDLE.

Conditional Generative Adversarial Networks (cGANs) can be used to generate new data from a specific labelled class of the training data (biologically-relevant, biologically-nonrelevant). In REKINDLE, we use cGANs to conditionally generate biologically relevant kinetic parameter sets using the biologically-relevant class label in the generator seed. While we are not interested in biologically-nonrelevant models because they do not represent any physiologically observed metabolic characteristics, for the sake of completion, we investigated can we generate non-relevant models using the corresponding class label in REKINDLE. We selected the generators with the highest incidence of biologically relevant models (Table S2) and generated 1000 non-relevant models for each metabolic physiology by changing the class label in the generator seed. We then tested these models for non-relevance by calculating the eigenvalues of their Jacobian matrix (Methods). The results are summarized in the Table below (Table S2).

**Supplementary Table S2:** Incidence of biologically relevant and non-relevant models generated with REKINDLE for four physiologies.

| | Physiology1 | Physiology 2 | Physiology 3 | Physiology 4 |
|---|---|---|---|---|
| Directionality of Reactions | $\underset{\rightarrow}{TALA}\ \underset{\rightarrow}{ICL}$ | $\underset{\rightarrow}{TALA}\ \underset{\leftarrow}{ICL}$ | $\underset{\leftarrow}{TALA}\ \underset{\rightarrow}{ICL}$ | $\underset{\leftarrow}{TALA}\ \underset{\leftarrow}{ICL}$ |
| Relevant Models | 97.7% | 97.3% | 99.3% | 100.0% |
| Non-relevant Models | 18.5% | 9.8% | 54.5% | 16% |
| (stable but slow, unstable) | (6.1%, 12.4%) | (3%, 6.8%) | (30.2%, 24.3%) | (11.1%, 4.9%) |

We observed that the incidence of non-relevant models is low compared to relevant models. This suggests that the cGANs did not learn the non-relevant space of the kinetic parameter space as well as it did for the relevant space. To investigate this space further, we discretized the kinetic parameter space of the training data into 16 subspaces based on the maximum eigenvalue of the Jacobian near the boundary line of our imposed classes i.e., $Re(\lambda_{max}) = -9,$ for all 4 physiologies (X axis, Supplementary Fig 12 a-d). For each of these 16 subspaces, we quantified the difference between the distributions of individual kinetic parameters that belong to a particular subspace and those subspaces that have larger eigenvalues, again using the KL divergence score. We observed that only locally stable subspaces ($Re(\lambda_{max}) < 0$) contained kinetic parameter sets that had significant differences in the distributions of certain kinetic parameters whereas non-relevant and especially locally unstable models ($Re(\lambda_{max}) > 0$) did not (Supplementary Fig 12 a-d, here the KL divergence scores are normalized for each subspace). This suggests that there are no determining characteristic parameters that make a certain kinetic parameter set locally unstable, unlike relevant parameter sets.

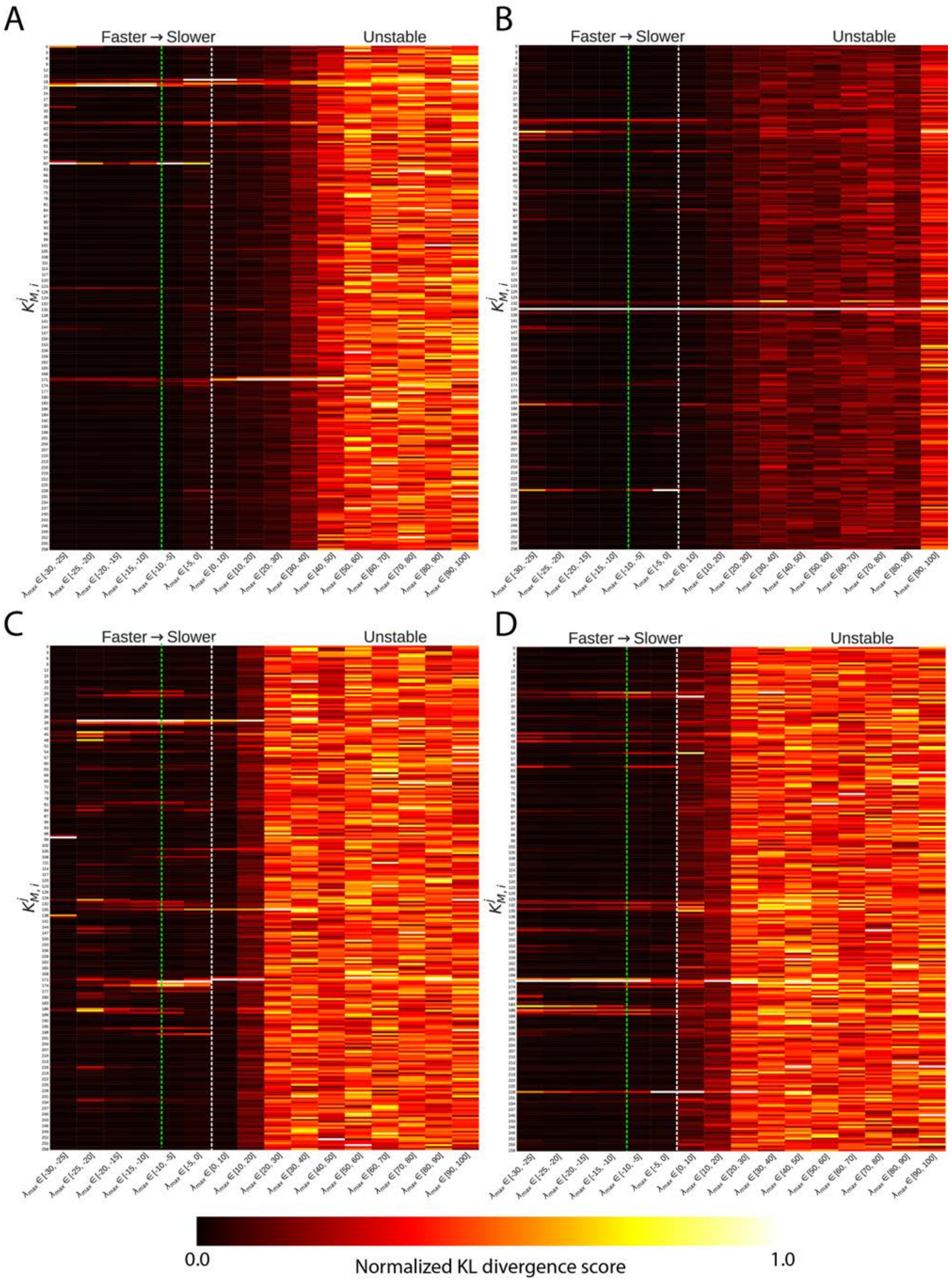

**Supplementary Figure 12:** Differences in distributions of individual kinetic parameters (y-axis) using KL divergence between 16 discretized sub-populations based on eigenvalue space (x-axis) near the class boundary i.e., $Re(\lambda_{max}) = -9$ (green dashed line) for (a) Physiology 1 (b) Physiology 2 (c) Physiology 3 (d) Physiology 4. The white dashed line represents $Re(\lambda_{max}) = 0$. Here the KL divergence for each sub-population is normalized between 0 and 1.

Moreover, we also plotted the same heatmap as Supplementary Fig. 12 but this time the KL divergence scores were normalized over the entire space and not for each subspace, to visualize where the strongest information was present. We observed that the strongest signals in the KL divergence scores of the relevant models appear further away from the eigenvalue partition line ($Re(\lambda_{max}) = -9$) for 3 of the 4 physiologies ( $Re(\lambda_{max}) \sim -20$ for physiology 1, $Re(\lambda_{max}) \sim -25$ for physiology 3 and $Re(\lambda_{max}) \sim -25$ for physiology 4 (Supplementary Figure 13 a-d). We hypothesize that the GANs used in REKINDLE is able to capture this bias and thus generates kinetic parameter sets that are further away from the eigenvalue class partition line compared to the training data (Figure 2e, Supplementary Fig 3a-c). Exceptionally, the strongest signal for physiology 2 occurs around $Re(\lambda_{max}) = 10$ (Supplementary Fig 13b, Supplementary Fig 13b).

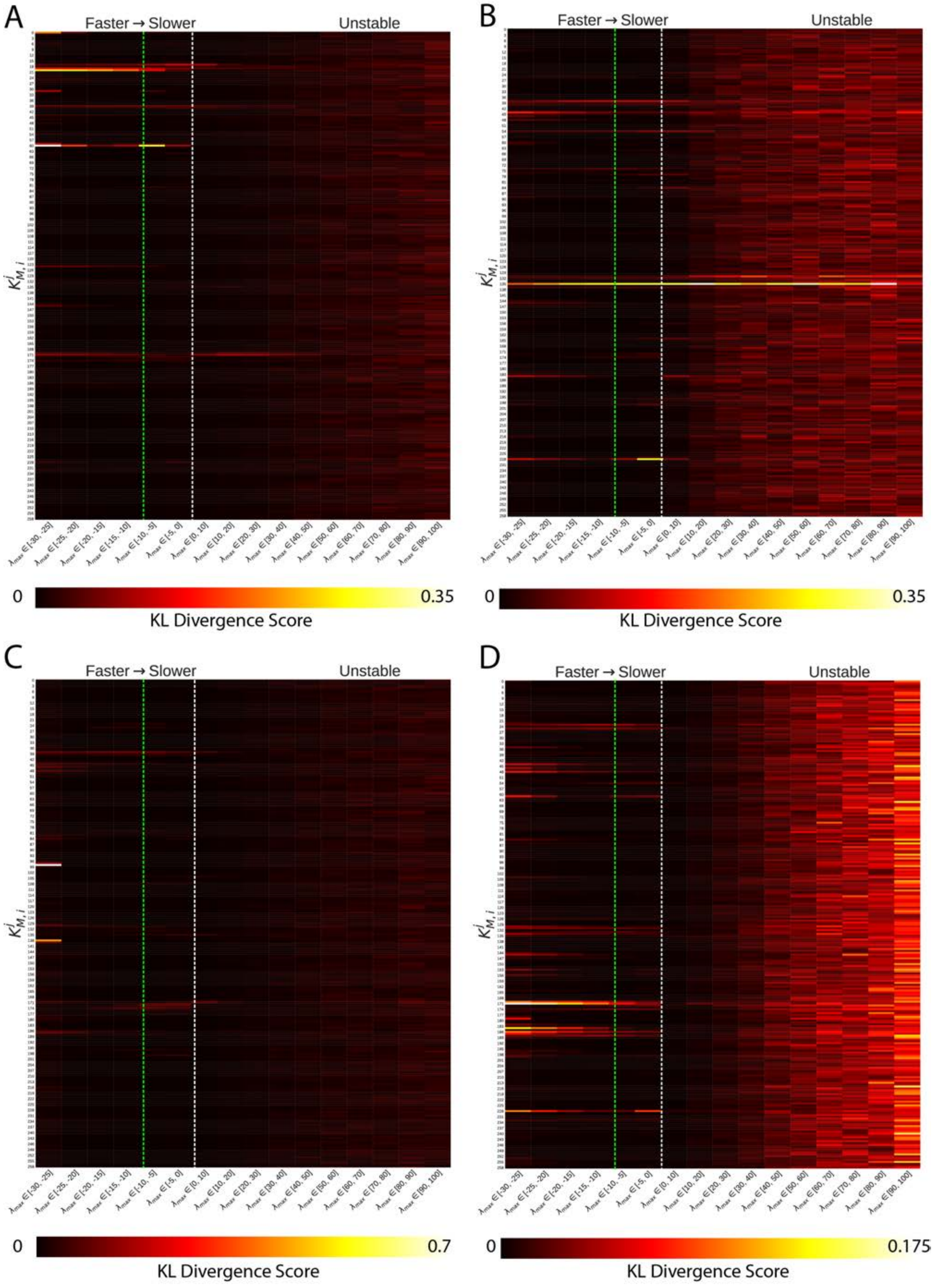

**Supplementary Figure 13:** Differences in distributions of individual kinetic parameters (y-axis) using KL divergence between 16 discretized sub-populations based on eigenvalue space (x-axis) near the class boundary i.e., $Re(\lambda_{max}) = -9$ (green dashed line) for (a) Physiology 1 (b) Physiology 2 (c) Physiology 3 (d) Physiology 4. The white dashed line represents $Re(\lambda_{max}) = 0$.

1. King, Z. A. et al. S1-Escher: A Web Application for Building, Sharing, and Embedding Data-Rich Visualizations of Biological Pathways. Plos Comput Biol 11, e1004321 (2015).
